# Membrane-gated SNARE zippering focuses energy for fusion

**DOI:** 10.64898/2026.08.16.745094

**Authors:** Avinash Kumar, Jie Yang, Lucas Anmolsingh, Anna R. Eitel, Zhiqun Xi, Lanxi Lin, Heidi E. Hamm, Yongli Zhang

## Abstract

Soluble N-ethylmaleimide–sensitive factor attachment protein receptors (SNAREs) drive stagewise membrane fusion by zippering into membrane-bridging four-helix bundles. Yet the conformations underlying successive fusion stages and the coupling of folding energy to bilayer remodeling remain unclear. Using optical tweezers, we measured the intermediates, energetics, kinetics, and force dependence of individual synaptic SNARE complexes assembled in cis on single membranes and in trans between apposed membranes. Membrane-anchored cis-SNAREs assembled through N-terminal and cooperative C-terminal/linker-domain transitions, whereas their transmembrane domains showed little intrinsic dimerization. Syntaxin retained membrane-dependent helical continuity through its linker domain before zippering was complete. PIP₂ strengthened but slowed late zippering. In trans, membrane repulsion arrested single trans-SNARE complexes in a half-zippered state. Gβγ further clamped this intermediate and inhibited late zippering; Gα–GDP, but not Gα–GTPγS, relieved the clamp, revealing a nucleotide-dependent mechanism for GPCR-mediated inhibition of neurotransmitter release. Modeling suggests that cooperative late zippering, syntaxin linker helicity, and concerted action of multiple SNAREs focus folding energy released over a long distance onto short-range membrane apposition. Thus, mechanically gated SNARE zippering is regulated by membrane forces, lipids, and regulatory proteins.

## INTRODUCTION

Neurotransmitter release requires synaptic vesicles to fuse with the presynaptic plasma membrane within milliseconds after an action potential. This reaction proceeds through rapid, tightly regulated transitions among distinct membrane and protein states, including docking, priming, fusion-pore opening, pore flickering or reclosure, and pore expansion (Figure 1A)^1–3^. The core fusion engine comprises the v-SNARE VAMP2/synaptobrevin on synaptic vesicles and the t-SNAREs syntaxin-1 and SNAP-25 on the plasma membrane; their coupled folding and assembly drive membrane fusion. The SNARE assembly is assisted by Munc18-1, Munc13-1, synaptotagmin (Syt), and other conserved regulators of neurotransmission^4–6^, and is further modulated by G protein-coupled receptors (GPCRs) and heterotrimeric G proteins^7^. Genetic and functional defects in the SNARE-dependent release machinery contribute to neurological disorders, diabetes, cancer, and immune disorders.^8^ Adrenergic GPCRs, in particular, regulate blood pressure, heart rate, and stress responses and are major targets of widely used drugs.^9^ Despite this broad physiological and clinical importance, how SNARE assembly drives successive stages of membrane fusion remains unclear^2^.

**Figure 1.**
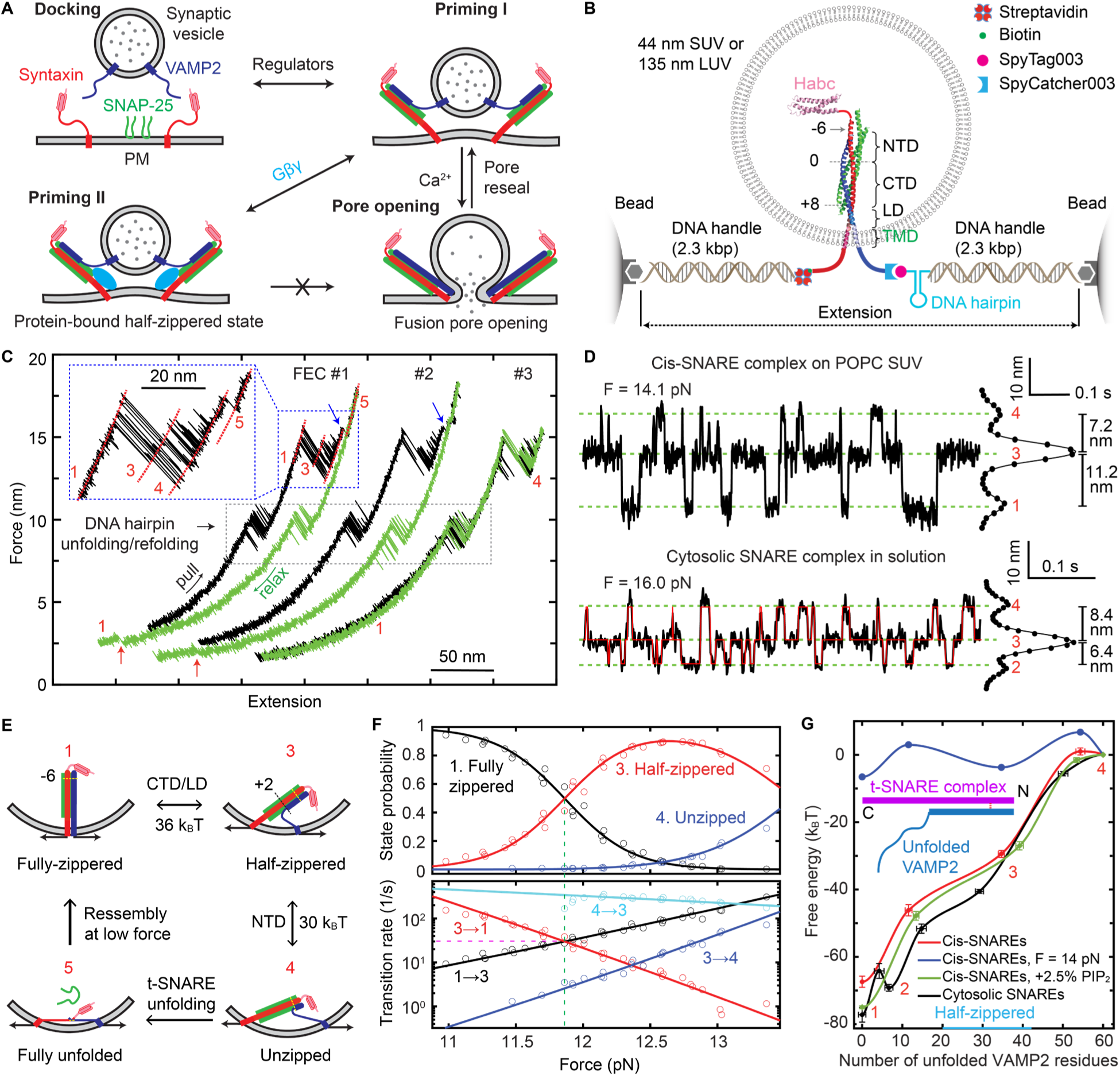
Membrane anchoring reshapes the folding-energy landscape of the synaptic cis-SNARE complex measured by optical tweezers. (A) Working model for Ca²⁺-triggered synaptic vesicle fusion and associated SNARE conformations. SNAREs and regulators dock vesicles near the plasma membrane (PM) and prime a partially zippered trans-SNARE state. Ca²⁺ triggers further SNARE zippering, which promotes fusion-pore opening and expansion. Gβγ stabilizes the partially zippered state by blocking further zippering. During kiss-and-run, the pore reseals, the SNAREs disassemble, and the vesicle eventually detaches from the PM. (B) Single-molecule optical-tweezers assay of a cis-SNARE complex embedded in an SUV or LUV. The fully assembled complex (PDB ID: 3HD7) was linked to two optically trapped beads through DNA handles and covalent or affinity linkages; an optional DNA hairpin served as an internal mechanical reference. NTD, CTD, LD, TMD, the syntaxin Habc domain, and SNARE layers −6, 0, and +8 are indicated. (C) Representative force-extension curves (FECs) from repeated pulling (black) and relaxation (green) of one cis-SNARE complex. FECs #1 to #3 are horizontally offset for clarity. Dashed boxes mark reversible DNA-hairpin unfolding/refolding and SNARE zippering/unzipping. Red state numbers denote the conformations in (E); red and blue arrows mark SNARE reassembly and t-SNARE unfolding, respectively. (D) Representative constant-mean-force extension trajectories for a cis-SNARE complex on POPC SUVs (top) and a cytosolic complex lacking TMDs (bottom). Green dashed lines mark mean state extensions; red lines show hidden Markov idealizations. Data were filtered to 1000 Hz. Right, extension histograms and corresponding extension changes. The structural interpretation of LD-unzipped state 2 is shown in Figure S7A. (E) Structural assignments of the observed states. CTD/LD zippering converts half-zippered state 3 to fully zippered state 1; NTD unzipping generates unzipped state 4, and t-SNARE unfolding generates fully unfolded state 5. Refolding at low force restores fully zippered state 1. (F) Force-dependent state probabilities (top) and transition rates (bottom). Symbols show measurements; solid lines show model fits. Green and magenta dashed lines mark the equilibrium force and rate of the CTD/LD transition, respectively. (G) Free-energy landscapes versus the number of VAMP2 residues unfolded from the C terminus for cis-SNARE complexes on POPC SUVs at zero force (red) or 14 pN (blue), on POPC/PIP₂ SUVs at zero force (green), and for the cytosolic complex at zero force (black). Symbols show measurements; solid lines are piecewise cubic fits. The unzipped state defines zero energy. State numbers mark energy minima; inset, unfolded VAMP2 residues relative to the t-SNARE template.

The prevailing zippering model proposes that membrane-bridging trans-SNARE complexes assemble from their membrane-distal N termini toward their membrane-proximal C termini, thereby drawing the vesicle and plasma membranes together (Figure 1A)^4,10,11^. Single-molecule and related studies have shown that zippering proceeds through discrete intermediates, including N-terminal-domain (NTD) assembly, a force-dependent half-zippered state, C-terminal-domain (CTD) zippering, linker-domain (LD) zippering, and possible final dimerization of the transmembrane domains (TMDs) (Figure 1B)^8,12–15^. These studies defined the energetics, kinetics, and force dependence of soluble SNARE assembly, but how SNAREs zipper in membranes remains poorly understood.

Three major gaps limit mechanistic understanding of SNARE-mediated membrane fusion. First, the energetics, intermediates, and kinetics of membrane-anchored SNARE zippering remain poorly defined, including their dependence on lipids and membrane geometry^13,16^. In particular, the roles of LDs and TMDs are unresolved: they may promote fusion through dimerization, lipid perturbation, force transmission, or some combination of these mechanisms^13,17–20^. In detergent, the fully assembled SNARE complex forms a parallel four-helix bundle, with continuous syntaxin and VAMP2 helices extending through their LDs and TMDs^13^. Truncating either or both TMDs reduced the thermal stability of the complex, consistent with stabilizing interactions involving the two TMDs. By contrast, replacing the VAMP2 TMD with polyvaline did not impair neurotransmission, and solid-state NMR revealed that the VAMP2 TMD is highly dynamic in lipid bilayers^21,22^. Thus, the extent and functional importance of TMD dimerization remain uncertain. The prefusion LD conformations are also unclear, with both folded and unfolded states proposed^5,8,18,23^. Rigid LDs have been hypothesized to transmit SNARE zippering forces to membranes^6,18,24^, but this hypothesis remains to be tested experimentally.

Second, how SNARE zippering is energetically coupled to membrane apposition and fusion remains poorly understood. SNARE zippering can begin when membranes are separated up to 20 nm and generate forces exceeding 15 pN^12,25^, whereas the primary opposing force—short-range membrane dehydration—decays exponentially with a decay length of ∼1 nm^26,27^. Much of the SNARE zippering reaction may therefore proceed against little resistance, dissipating the zippering energy before the membranes are close enough to fuse^20,28,29^. This potential mismatch is particularly relevant to CTD zippering, which, based on measurements of cytosolic SNARE complexes, would occur at membrane separations greater than 2.5 nm with unfolded LDs^25,30^. Alternative models propose that LDs promote fusion either by locally reorganizing lipids in a detergent-like manner^20^ or, when unfolded, by acting as entropic springs that draw the two membranes together^29^. In both models, CTD zippering primarily brings the membranes into proximity, leaving its substantial energy poorly coupled to the short-range fusion barrier.

Third, how regulators act on specific SNARE assembly states remains difficult to dissect^5,8^. Many regulators also bind membranes, and short-range membrane forces may alter the intermediates and kinetics measured for cytosolic SNAREs under approximately constant force^12,14^, necessitating measurements in a membrane context. This issue is particularly relevant to regulation of neurotransmitter release through presynaptic Gi/o-coupled GPCRs, including α2A adrenergic receptors, 5-HT1B serotonin receptors, GABA_B receptors, and µ-opioid receptors^9^. Their activation enables membrane-anchored Gβγ to bind directly to the SNARE machinery and inhibit neurotransmitter release^7,31^. The molecular mechanism by which Gβγ regulates SNARE assembly remains poorly understood. Because SNAREs readily form off-pathway misassembled complexes that confound ensemble measurements, addressing these questions requires single-molecule measurements in native-like membranes with high spatiotemporal resolution and controlled mechanical load to distinguish correctly assembled SNARE complexes from misassembled species^8,14,15^.

## RESULTS

### Reversible cis-SNARE complex assembly on SUV membranes

Optical tweezers have rarely been used to quantify membrane protein folding and dynamics^32^. We reconstituted single purified cis-SNARE complexes on vesicles and linked the C termini of syntaxin and VAMP2 to optically trapped beads through two DNA handles^33^ (Figure 1B and Figures S1 to S3). This geometry captured nearly the full zippering energy, minimized membrane-imposed opposing load, and enabled comparison of cytosolic, cis-, and trans-SNARE assemblies. Magnetic tweezers were previously used to monitor GlpG folding in vesicles and bicelles, but quantitative energetics and detailed kinetics were derived in bicelles rather than vesicles^34^. We pulled and relaxed individual complexes parallel to the vesicle membrane at 10 nm s^-1^ and inferred conformational changes from tether extension and force. An optional DNA hairpin in series with VAMP2 served as a single-molecule marker. To test curvature effects, we used small and large unilamellar vesicles (SUVs and LUVs) with diameters of 44 ± 13 and 135 ± 36 nm, respectively (mean ± SD; Figure S4). To permit repeated reassembly and equilibrium measurements, syntaxin and VAMP2 were crosslinked by a disulfide bond at the N-terminal −6 layer in most experiments^25^.

Repeated pulling and relaxation of single cis-SNARE complexes on POPC SUVs produced force-extension curves (FECs; Figure 1C). The DNA-hairpin transition at ∼10 pN confirmed that single VAMP2 molecules were pulled. The transition near 14 pN reflected three-state SNARE folding and unfolding (Figure 1C, inset), resembling the CTD and NTD transitions of cytosolic complexes^25^. A small irreversible extension jump at 17.1 ± 2.7 pN, followed by no further unfolding, was consistent with complete t-SNARE unfolding (Figure 1C)^12^. Relaxation to low force reassembled the complex, yielding nearly identical unfolding patterns in subsequent cycles (FEC #2). Reversing the pull before t-SNARE unfolding instead produced overlapping pulling and relaxation FECs (FEC #3), indicating rapid, reversible zippering on an intact t-SNARE template. We detected no distinct LD or TMD transitions, even without the DNA hairpin (Figure S5), suggesting that membrane anchoring changes the cis-SNARE zippering pathway.

### Membranes reshape the energy landscapes of SNARE assembly

To resolve SNARE transitions at higher spatiotemporal resolution, we held single cis-SNARE complexes at different trap separations and measured equilibrium transitions among three states (Figure 1D and Figure S6). The most extended state (state 4) was assigned to the unzipped complex, because it resembled the unzipped state of cytosolic complexes (Figure 1D; Figure 2A, compare POPC-SUV with Cyto; Figure S7)^25^, preceded t-SNARE unfolding, and disappeared without the N-terminal crosslink (Figure 1C and Figure S4). States 3 and 1 were assigned to half-zippered and zippered SNAREs, respectively (Figure 1E). Thus, membrane-anchored cis-SNARE complexes assemble stepwise, like cytosolic complexes^12,30^.

**Figure 2.**
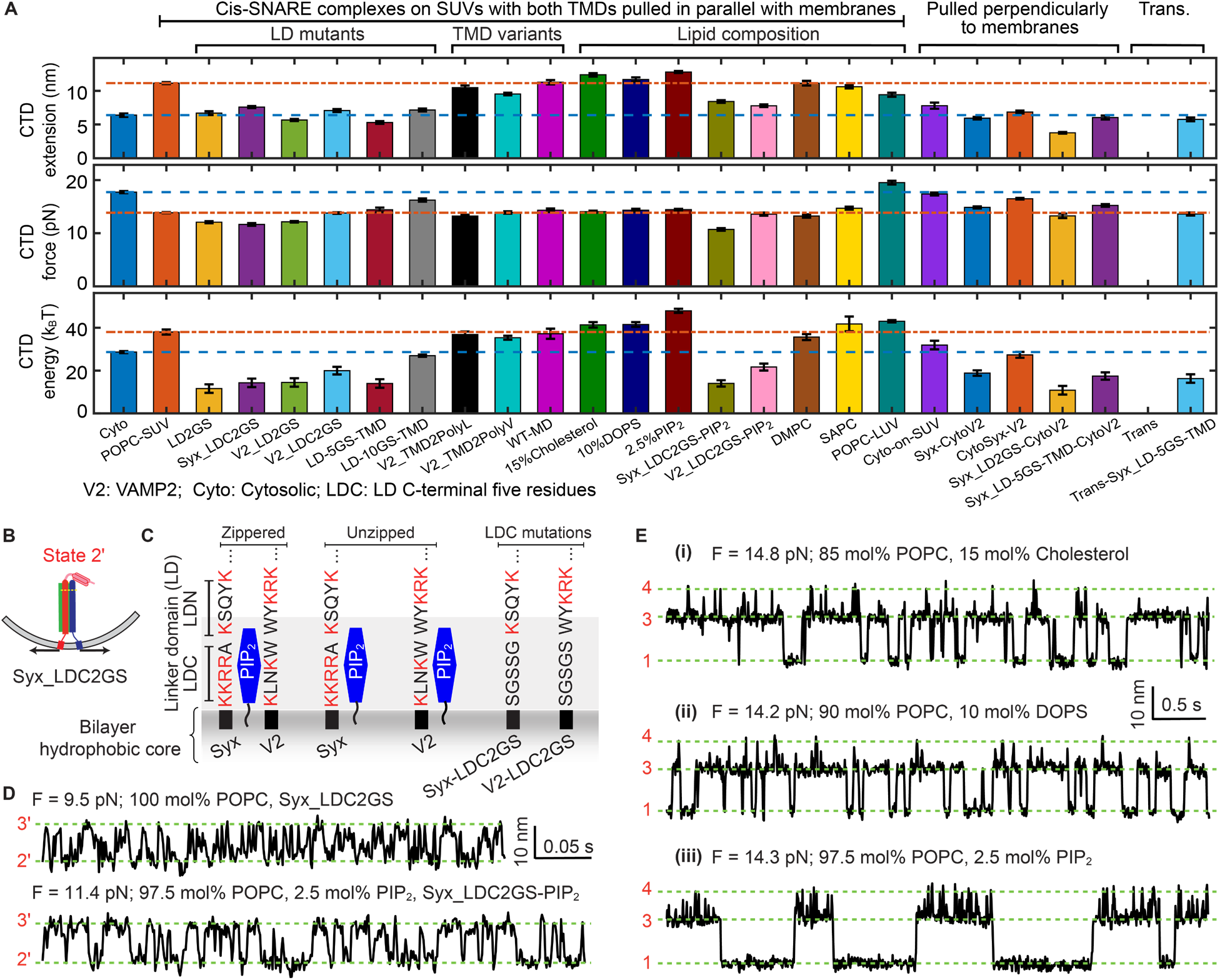
Effects of SNARE modifications and lipids on cis-SNARE complex assembly. (A) Mean extension changes, equilibrium forces, and free energies for CTD/LD or CTD transitions of the indicated WT or mutant SNARE complexes, lipid compositions, bilayer thicknesses, and membrane-anchoring configurations. Blue and red dashed lines mark values for the cytosolic complex and WT cis-SNARE complex on POPC SUVs, respectively. Bars and error bars show mean ± SEM. No CTD transition was detected for the trans-SNARE complex (Trans). NTD parameters are in Figure S10. See Table S1 for sample details. (B) Schematic of the LD-unfolded state in the mutant cis-SNARE complex. (C) Syntaxin (Syx) and VAMP2 (V2) LD sequences used to test electrostatic coupling to PIP₂-containing membranes. Basic residues proposed to contact PIP₂ are highlighted in red; glycine-serine substitutions identify the corresponding LD mutants. (D) Representative constant-force trajectories of the Syx_LDC2GS cis-SNARE complex on POPC SUVs or SUVs containing 97.5 mol% POPC and 2.5 mol% PIP₂. Scale bars are indicated. Data were filtered to 1000 Hz. (E) Representative extension-time trajectories of WT cis-SNARE complexes at the indicated mean forces on SUVs containing 15 mol% cholesterol (i), 10 mol% DOPS (ii), or 2.5 mol% PIP₂ (iii). Green dashed lines mark assigned states defined in Figure 1E. Data were filtered to 500 Hz.

Membrane anchoring and lipid bilayers reshaped the SNARE energy landscape. First, the extension change between zippered and half-zippered states was larger for cis-SNARE than cytosolic complexes [11.1 ± 1.8 nm (N = 163) versus 6.4 ± 1.7 nm (N = 47); Figures 1D and 2A], suggesting that an extended C-terminal region, likely including LDs, participates in zippering. Replacing the entire LD or its C-terminal half in syntaxin (Figures 2B and 2C, Syx_LDC2GS; Table S1), VAMP2 (V2_LD2GS and V2_LDC2GS), or both (LD2GS) with flexible glycine-serine (GS) linkers impaired this transition (Figure 2D, top trace; Figure 2A), indicating that the CTDs and LDs form a cooperative CTD/LD folding domain. Inserting 5-or 10-residue GS linkers between the LDs and TMDs also weakened CTD/LD zippering (Figure 2A, LD-5GS-TMD and LD-10GS-TMD), indicating that TMD proximity stabilizes late zippering. This stabilization is consistent with helical continuity from the LDs into the TMDs in the fully zippered complex^13^, but not with strong intrinsic TMD dimerization^21,22^, Neither replacing the VAMP2 TMD with polyleucine or polyvaline sequences (Figure 2A, V2_TMD2PolyL and V2_TMD2PolyV) nor restoring native cysteines in the syntaxin and VAMP2 TMDs (WT-TMD) measurably altered SNARE assembly (Figures S8 and S9)^21^. LD and TMD mutations had little effect on NTD transitions (Figure S10). Second, cis-SNARE complexes zippered at lower forces than cytosolic complexes (Figure 2A and Figure S7C); solubilizing the same complex in 2% n-octyl-β-D-glucopyranoside (OG) shifted the transition to higher force (Figure S11), showing that this effect was bilayer-dependent. Finally, the CTD/LD equilibrium transition rate was lower than the cytosolic CTD rate (18 ± 4 versus 86 ± 3 s^-1^, mean ± SEM; Figure 1F), whereas NTD rates were similar. Thus, bilayers merge the LD and CTD into one cooperative transition without detectable TMD dimerization.

Hidden Markov analysis yielded state probabilities, forces, extensions, and transition rates as functions of mean force^35^ (Figure 1F and Figure S6). A force-dependent folding model converted these measurements into simplified energy landscapes at zero force and under load (Figures 1F and 1G; Figure S12)^36^. Without force, cis-SNARE folding was downhill (Figure 1G), indicating that the half-zippered intermediate and stepwise zippering are populated only under biologically relevant opposing forces. Cis-SNARE zippering released less energy than cytosolic-complex zippering (67 ± 2 versus 77 ± 2 k_B_T). This difference was not simply due to TMD confinement, because long GS linkers between the LDs and TMDs restored CTD and NTD zippering energies to near cytosolic values (Figure 2A, compare LD-10GS-TMD with Cyto). The cis-SNARE half-zippered state also shifted from the +3 toward the +1 layer, potentially lengthening the fusion power stroke. Because this intermediate is structurally plastic during force-induced unfolding, we classified states with 20 to 43 unfolded VAMP2 residues—approximately layers −1 to +6—as half-zippered (Figure 1G).

We reexamined cytosolic SNARE complexes using otherwise identical constructs lacking TMDs (Figures 1D and S7B). CTD and NTD energies and kinetics matched previous measurements^12,25^. LD substitutions showed that the syntaxin LD and N-terminal VAMP2 LD are structured in the half-zippered state, whereas the C-terminal VAMP2 LD folds and unfolds asymmetrically on a structured syntaxin LD (Figure S7A). This revises our earlier view that both LDs become disordered in the LD-unzipped state. The results support the t-SNARE template used to derive the cis-SNARE energy landscape and imply that syntaxin forms a continuous helix from its CTD through the LD to the TMD in the half-zippered state (Figure 1E), consistent with a large tilt relative to the membrane plane^16,24^. Persistence of the folded syntaxin LD in the unzipped state also explains the lower total zippering energy of cis-SNARE than cytosolic complexes (compare state 4 in Figure 1E with Figure S7A).

### PIP₂ strengthens but slows SNARE zippering

To test lipid effects, we incorporated fusion-modulating lipids into SUVs^37,38^. Cholesterol (15 mol%) or DOPS (10 mol%) only slightly increased CTD/LD zippering extension and energy and minimally affected NTD zippering (Figure 2E, i and ii; Figure 2A, 15%Cholesterol and 10%DOPS; Figure S10). By contrast, 2.5 mol% PIP₂ increased CTD/LD extension and energy (Figure 2A, 2.5%PIP₂) and slowed CTD/LD zippering ∼10-fold without significantly changing the NTD rate (Figure 2E, iii; Figure S13). PIP₂ may stabilize folded CTD/LD states through multivalent electrostatic interactions with lysine and arginine residues in the syntaxin and VAMP2 LDs, and interactions with unfolded LDs may slow zippering (Figure 2C). Consistently, PIP₂ had little effect when the C-terminal half of either LD was replaced by GS linkers (Figure 2D, bottom trace). Replacing POPC with shorter-chain DMPC or longer-chain SAPC did not significantly alter SNARE zippering (Figure 2A, DMPC and SAPC; Figure S10), suggesting that flexible TMDs accommodate hydrophobic mismatch^21,22^. Thus, among the lipids tested, PIP₂ selectively and strongly altered late zippering energetics and kinetics.

### Membrane geometry modulates SNARE zippering through syntaxin linker bending

To test whether membrane curvature alters SNARE zippering^39^, we examined cis-SNARE assembly on 135-nm LUVs (Figure 3A). Relative to SUVs, LUVs shifted both transitions to higher forces and energies (Figure 3B; Figure 3C, #1; Figure 2A, POPC-LUV; Figure S10). CTD/LD zippering on LUVs had a higher equilibrium force (19.5 versus 13.9 pN), smaller extension change (9.4 versus 11.1 nm), and higher free energy (43.1 versus 38.0 k_B_T); total zippering energy was 11.4 k_B_T greater (Figure S14). We infer that, on the flatter LUV membrane, unzipping partially or completely unfolds the syntaxin LD in the half-zippered and unzipped states (Figure 3A). Assuming the same fully zippered state on SUVs and LUVs, we attribute the 11.4 k_B_T difference to syntaxin LD unfolding in their distinct unzipped states (Figure 3A, state 4′, versus Figure 1E, state 4).

**Figure 3.**
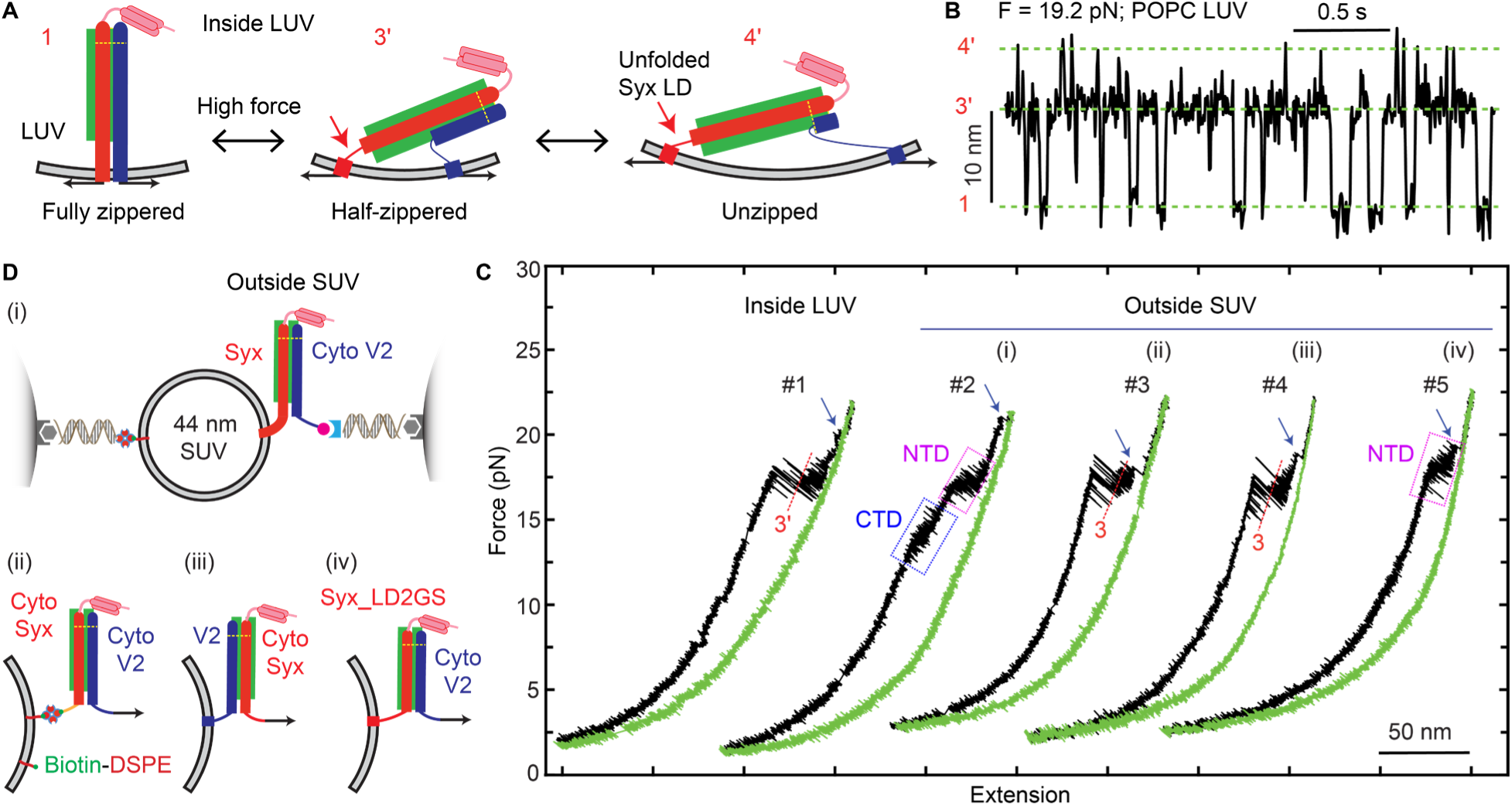
Membrane curvature and pulling geometry alter SNARE assembly. (A) Model for the CTD/LD transition on an LUV. Sharp bending of the syntaxin linker domain (Syx LD) in the half-zippered and unzipped states promotes LD unfolding (red arrow). (B) Representative extension-time trajectory of a WT cis-SNARE complex on a POPC LUV at the indicated mean force. Green dashed lines mark discrete folding states. Data were filtered to 200 Hz. (C) Representative force-extension curves (FECs) for single SNARE complexes in the indicated LUV and SUV configurations. Pulling geometries are schematized in (A), #1, and (D), #2 to #5. Pulling and relaxation traces are shown in black and green, respectively. Blue arrows mark SNAP-25 dissociation after t-SNARE unfolding. Blue and magenta dashed boxes mark CTD and NTD transitions, respectively. (D) Pulling geometries for complexes anchored to SUVs through native syntaxin or VAMP2 TMDs (i, iii, and iv) or engineered flexible linkers (ii). Arrows show force direction.

We next pulled single SNARE complexes perpendicular to SUV membranes (Figure 3D, i), anchoring each through either the syntaxin or VAMP2 TMD and pulling from the partner lacking its TMD. SUVs withstood the unfolding force without substantial deformation, as cytosolic SNARE measurements were nearly identical with or without SUVs (Figure 3D, ii; Figure 3C, #3; Figure 2A, compare Cyto-on-SUV with Cyto). Anchoring through the syntaxin TMD lowered CTD/LD force and energy, whereas anchoring through the VAMP2 TMD had little effect (Figure 3D, i and iii; Figure 3C, #2 and #4; Figure 2A, Syx-CytoV2 and CytoSyx-V2), indicating that force-induced syntaxin-LD bending destabilizes CTD/LD zippering. Consistently, replacing the syntaxin LD with a GS linker or inserting a five-residue GS linker between its LD and TMD strongly destabilized the transition (Figure 2D; Figure 2A, Syx_LD2GS-CytoV2 and Syx_LD-5GS-TMD-CytoV2). Replacing the syntaxin LD eliminated the transition in two-thirds of FECs (Figure 3C, #5), whereas NTD zippering was minimally affected (Figure S10). Thus, before late zippering, membrane geometry modulates assembly primarily through helical continuity and bending of the syntaxin LD, whereas the VAMP2 LD is more flexible.

### Single trans-SNARE complexes are half-zippered under membrane load

To examine SNARE complexes bridging two membranes, we attached t-and v-SNAREs to 44-nm SUVs and ∼13-nm nanodiscs, respectively, and formed trans-SNARE complexes crosslinked at the −6-layer (Figure 4A; Figure S15; Method details). Previous fusion assays have used defined average numbers of trans-SNARE complexes between nanodiscs and SUVs or free-standing planar lipid bilayers^40,41^. These studies reported that one complex can induce lipid mixing, whereas three are required to dilate a fusion pore. In our assay, single trans-SNARE complexes were tethered between beads and pulled through one VAMP2 molecule per nanodisc, as confirmed by the DNA-hairpin signature (Figure 4B). Pulling revealed reversible NTD unfolding and irreversible t-SNARE unfolding, but no discrete CTD/LD transition in 52 FECs from 35 complexes (Figure 4B, FECs #1 and #2; Figures 4C and 2A; Figure S10). Fully unfolded complexes reassembled at low force in 2 µM SNAP-25 but again showed only the NTD transition (Figure 4B, #2), indicating that fusion had not occurred. Inserting a five-residue GS linker between the syntaxin LD and TMD restored the CTD/LD transition in 18 of 19 experiments, although below the NTD transition force (Figure 4B, #3; Figure 2A, Trans-Syx_LD-5GS-TMD). The insertion thus increased membrane separation, weakened repulsion, and permitted further zippering (Figure 4A). We conclude that single trans-SNARE complexes predominantly remain half-zippered under membrane load.

**Figure 4.**
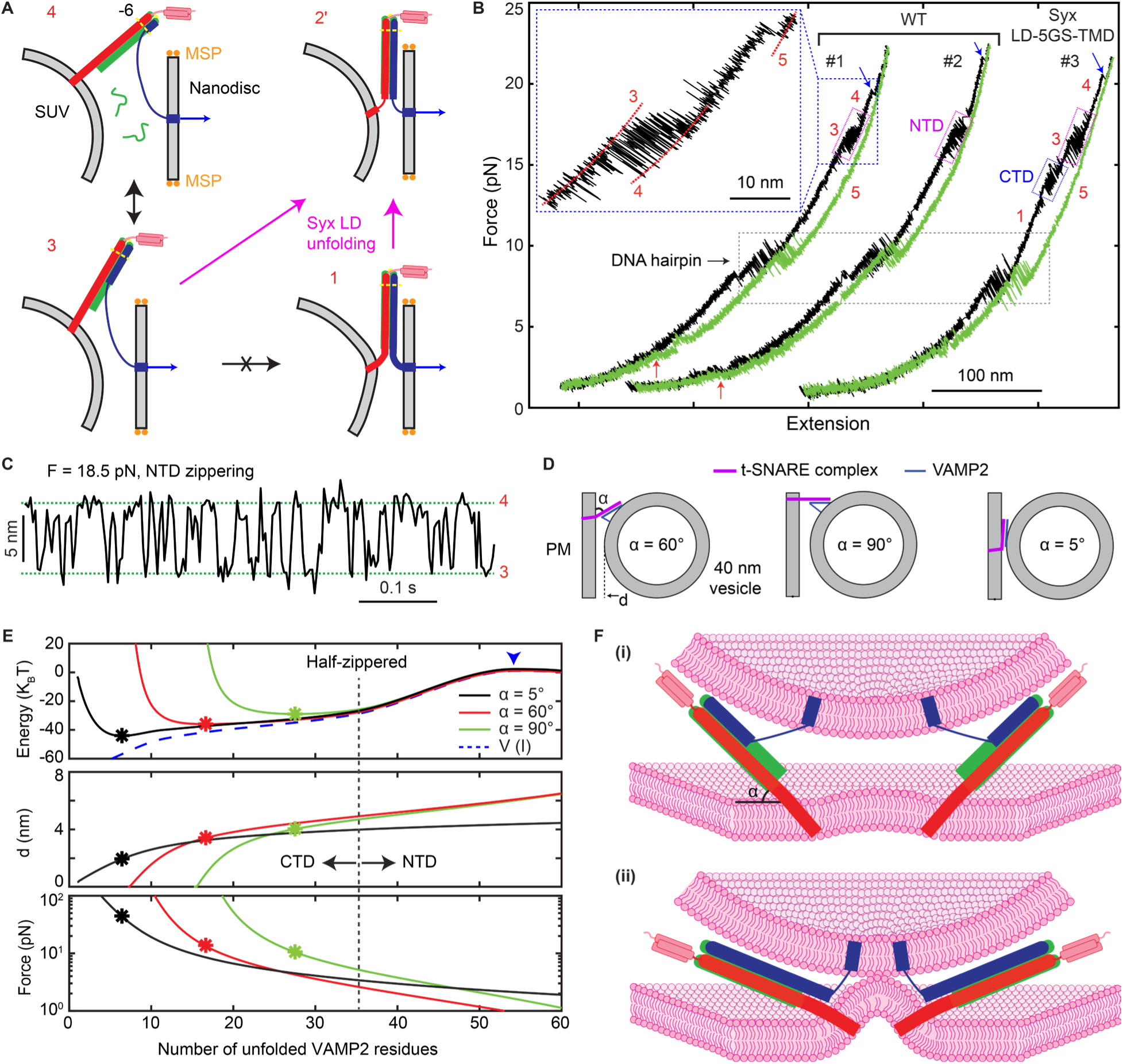
Single trans-SNARE complexes are half-zippered. (A) Optical-tweezers assay for single trans-SNARE complexes formed between t-SNARE-bearing SUVs and VAMP2-bearing nanodiscs. Representative SNARE states, membrane scaffold proteins (MSP), and pulling directions (blue arrows) are indicated. A nearly perpendicular t-SNARE complex stabilizes half-zippered state 3; membrane bending by additional SNARE complexes or syntaxin LD unfolding promotes zippering to state 1 or 2′, respectively. (B) Representative force-extension curves from pulling (black) and relaxing (green) individual WT trans-SNARE complexes or the Syx_LD-5GS-TMD mutant. Dashed boxes mark unfolding/refolding transitions of CTD (blue), NTD (magenta), and the DNA hairpin (black). Red and blue arrows mark SNARE reassembly and t-SNARE unfolding, respectively. State numbers denote the SNARE conformations schematized in (A) or defined in Figure 1E. (C) Representative constant-force extension trajectory showing reversible NTD zippering and unzipping between states 3 and 4 of one WT trans-SNARE complex. Green dashed lines mark the two conformational states. Data were filtered to 500 Hz. (D) Calculated equilibrium SNARE conformations and membrane separations at three tilt angles (á) of the helical t-SNARE template relative to the planar plasma membrane. The t-SNARE template and both membranes were treated as rigid. (E) Calculated minimum total free energy (top), equilibrium membrane separation (middle), and force generated by the trans-SNARE complex (bottom) versus unfolded VAMP2 residues for the indicated α values. Stars mark global minima; the blue arrowhead indicates the energy barrier for unzipping. The blue dashed curve is the intrinsic WT cis-SNARE folding energy from Figure 1G; the vertical black dashed line marks its half-zippered state. (F) Proposed energy-focusing model for SNARE-mediated membrane apposition. Cooperative zippering within and among trans-SNARE complexes, continuous syntaxin helicity, and membrane bending together draw membranes toward point contact. One or two complexes remain half-zippered (i); whereas additional SNAREs bend the membrane and promote further cooperative zippering (ii).

### SNARE assembly focuses energy onto membrane apposition

To explain the half-zippered trans-SNARE state and the coupling between SNARE zippering and membrane apposition, we simulated one or more complexes using the measured cis-SNARE folding landscape and a reported membrane-repulsion potential^26^ (Figure S16). The potential had an amplitude of 100 k_B_T and decayed exponentially with a decay distance of 0.8 nm (Supplementary Text, Sections T4 and T5). Because syntaxin tilt in a partially zippered complex is unknown, we modeled fixed angles of 5°, 60°, and 90° relative to the plasma-membrane plane (Figure 4D). Total free energy included intrinsic zippering energy, membrane repulsion, and entropic stretching of unfolded VAMP2^36^ and was minimized over VAMP2 unzipping and membrane separation (Figure S17A). At α = 60°, the global minimum contained 16.6 unfolded VAMP2 residues at a membrane separation of 3.4 nm (Figure S17A), consistent with reported docked-vesicle separations^3,42^.

Across tilt angles, NTD zippering encountered little membrane resistance and brought the membranes to separations below 6.5 nm with SNARE stretching forces below 5 pN (Figure 4E and Figure S17B). CTD/LD zippering, however, first lowered and then raised total energy as membrane resistance increased sharply, producing an equilibrium state with a partially zippered complex at separations > 2 nm. Predicted unzipped VAMP2 increased with syntaxin tilt: ∼6, ∼17, and ∼28 residues at α = 5°, 60°, and 90°, respectively, with corresponding increases in energy (Figure 4E). The measured 22 ± 2 unfolded residues suggest a tilt between 60° and 90°. This equilibrium occupied a shallow well only 3 k_B_T below the half-zippered cis-SNARE state on POPC SUVs; analysis versus membrane separation yielded the same conclusion (Figure S17C).

A more upright syntaxin orientation (α ≥ 60°) strongly enhanced coupling between zippering and membrane apposition. At α = 5°, the trans-SNARE complex was trapped behind a fusion barrier of ∼33 k_B_T even with a fully assembled SNARE complex and an unzipping barrier of ∼31 k_B_T at 54 unfolded VAMP2 residues (Figure 4E and Figure S18A), requiring many complexes to lower the calculated fusion barrier (Figure S18B). Large tilt angles instead allowed partially zippered complexes to store energy (Figure 4E; Figure 4F, i) that could bend the adjacent membrane to generate point contact or distort lipids to facilitate fusion (Figure 4F, ii)^18,27,42^. Three or four complexes eliminated the calculated fusion barrier at 60° or 90°, respectively (Figure S18B). The model helps explain why disrupting helical continuity in syntaxin or related Qa-SNAREs, but not VAMP2, impairs fusion^43–47^, why larger tilt angles facilitate fusion^16^, and why fusion depends on membrane curvature and bulky proteins bound to the SNARE bundle^39,48^. Cooperative CTD/LD folding and t-SNARE helical continuity therefore focus energy released over the long zippering distance onto short-range membrane apposition.

### Gβγ clamps half-zippered SNARE complexes

Activated GPCRs catalyze GDP-to-GTP exchange on Gα, promoting dissociation of heterotrimeric Gαβγ into Gα-GTP and Gβγ (Figure 5A). To test how Gβγ affects zippering, we reconstituted purified geranylgeranylated Gβγ with SNARE complexes into SUVs and pulled single complexes perpendicular to the membranes (Figure 5B and Figure S19). Gβγ produced long pauses near the half-zippered state during the rapid CTD/LD transition, thereby clamping half-zippered complexes (Figure 5C, i and ii). Dwell times ranged from tens of milliseconds to minutes, with survival time constants of 5.3 ± 0.1 s and 65 ± 5 s and an estimated clamping rate of ∼12 min^-1^ (Figures 5D and 5E). Pauses also occurred less frequently in the fully zippered state (1.8 ± 0.6 min^-1^; Figure 5C, iii), whereas Gβγ had little effect on NTD zippering or unzipping (Figure 5C, iv). Preferential clamping of the half-zippered state also occurred with cytosolic complexes and soluble Gβγ (Figure 5C, v; Figure S20). Transition kinetics between Gβγ-clamped half-zippered and unzipped states suggest that Gβγ remains associated during unzipping (Figure 5B, state 4; Figure S20), consistent with its reported binding primarily to the SNAP-25 C-terminus^31^. Thus, Gβγ preferentially clamps CTD/LD zippering, providing a direct mechanism for inhibiting neurotransmitter release.

**Figure 5.**
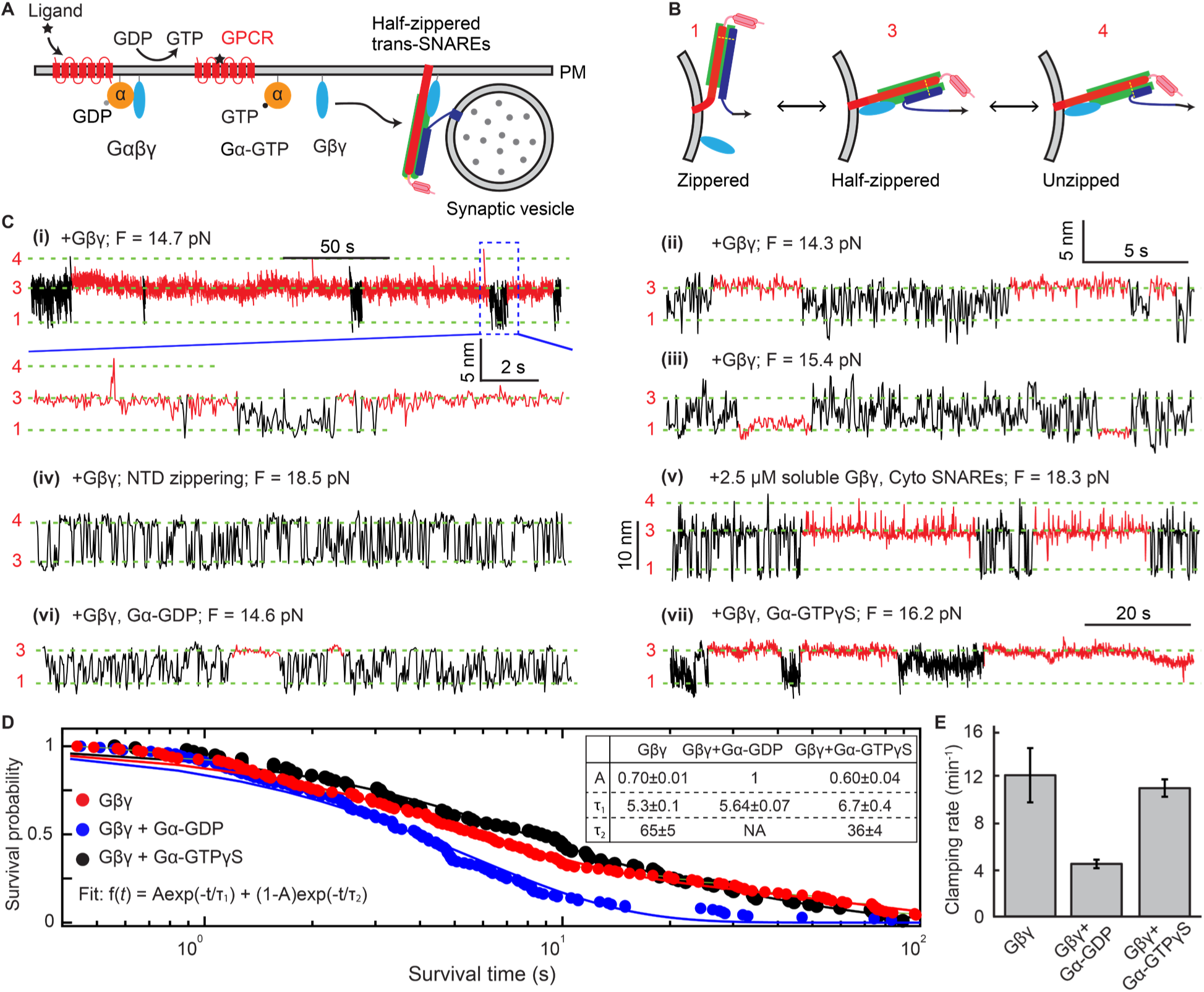
Gβγ clamps the half-zippered SNARE complex, and Gα–GDP attenuates this inhibition. (A) GPCRs inhibit neurotransmitter release by enabling Gβγ to block full SNARE zippering. (B) SNARE conformations and Gβγ-binding states underlying Gβγ-dependent inhibition of SNARE zippering. (C) Constant-force extension traces with Gâã alone (i to v), Gâã plus Gá**–**GDP (vi), or Gβγ plus Gα**–**GTPγS (vii). Gβγ caused long dwells in the zippered state (iii) or half-zippered state (other traces), highlighted in red, amid rapid SNARE folding and unfolding transitions (black). The boxed region in i is expanded below. Unless otherwise indicated, all traces use the scale bars in ii. Data were filtered to 50 Hz. Trace v is also shown at 1 kHz in Figure S20. (D) Survival probabilities of the Gâã-clamped half-zippered state versus dwell time without or with Gα**–**GDP or Gα**–**GTPγS. Measurements (dots) were fit with one or two exponentials (curves); fitting parameters are shown in the inset table. (E) Gâã-clamping rates for the half-zippered SNARE complex without or with Gα**–**GDP or Gα**–** GTPγS.

We next added Gα–GDP or Gα–GTPγS to test how the Gα nucleotide state modulates Gβγ-dependent clamping. Gα-GDP reduced the clamping rate ∼3-fold and shortened the clamped-state lifetime (Figure 5C, vi; Figures 5D and 5E). The survival probability fitted a single exponential with the shorter time constant, suggesting that Gα–GDP displaces Gβγ from the long-lived state and may rapidly relieve fusion inhibition. By contrast, Gα–GTPγS had little effect on clamping rate and only modestly shortened the mean clamped-state lifetime (23 s without versus 18 s with Gα–GTPγS; Figure 5C, vii; Figures 5D and 5E). Thus, Gβγ clamps CTD/LD zippering to inhibit fusion, whereas Gα–GDP, but not Gα–GTP, binds Gβγ to relieve the clamp and restore zippering (Figure 5A).

## DISCUSSION

We developed optical-tweezers assays to measure membrane protein folding on single membranes and between apposed membranes under controlled load. Applied to neuronal SNAREs, these assays show that membranes reshape the SNARE folding landscape rather than merely anchor SNAREs. In cis, SNAREs zipper through two principal steps: NTD assembly and cooperative CTD/LD assembly. The late transition depends on helical continuity in syntaxin from the CTD through the LD to the TMD, whereas the VAMP2 LD is more plastic. TMDs enhance CTD/LD zippering by stabilizing LD helices rather than through direct TMD dimerization^13^. PIP₂ stabilizes but slows CTD/LD zippering. In trans, membrane repulsion arrests single complexes in a half-zippered state. This finding seems at odds with earlier reports that a single SNARE complex can mediate lipid mixing and transient pore opening^40,41,49^, processes that likely require at least transient full zippering. These observations may be reconciled if a single half-zippered trans-SNARE complex undergoes rare excursions toward the fully zippered state during prolonged observations^50^, or if events statistically attributed to single complexes involved multiple complexes. Modeling suggests that cooperative CTD/LD folding, a continuous membrane-anchored syntaxin helix, and multiple SNARE complexes focus long-range folding energy onto short-range membrane apposition. Finally, Gβγ clamps the half-zippered state without affecting NTD zippering, indicating that it targets a late, partially zippered priming intermediate^51^ and directly mediates GPCR inhibition of neurotransmitter release. These findings define SNAREs as mechanically gated folding machines whose activity is tuned by membranes, lipids, and regulatory proteins to remodel membranes at the fusion site.

A balance between the driving force of SNARE zippering and opposing membrane repulsion may enable reversible fusion-pore dynamics, analogous to a seesaw poised near equilibrium^1,3,41^. The shallow energy minimum predicted for the trans-SNARE intermediate may facilitate rapid forward zippering for pore opening, reverse unzipping for pore closure during flickering or kiss-and-run, and regulation by proteins that bind SNARE intermediates weakly^5^. The tight coupling also enhances the energy efficiency of membrane fusion. The sensitivity of synaptic vesicle fusion to nearly all hydrophobic-layer mutations further supports tight coupling between zippering and fusion: even mutations that only modestly weaken CTD zippering markedly impair fusion^25,52–54^. Such sensitivity is difficult to reconcile with models in which fusion is driven predominantly by local detergent-like or entropic effects^20,29^. Instead, it strongly supports a primarily mechanical mechanism,^4^ while allowing lipid perturbation and other effects to contribute.

This work provides a framework for testing how membrane composition, geometry, and regulatory proteins tune SNARE-mediated fusion. Perturbations with modest effects in cis may act more strongly in trans, where a second membrane opposes zippering^55^. The syntaxin LD helix sustains axial forces up to 16 pN without its TMD (Figure 2A; Figure S7B, iii) but can fail near 14 pN under force parallel to the membrane (Figures 3C and 2A). The syntaxin tilt angle, extent of membrane bending, and their coupling to fusion remain uncertain. Future applications can test cooperative assembly of multiple trans-SNARE complexes and how other regulators alter assembly and fusion.

## ACKNOWLEDGMENTS

We thank Jim Rothman for discussions; David Zenisek, Chenxiang Lin, and Min Wu for manuscript review; and Erdem Karatekin, Fei Xu, Rebekah Jackson, and Diana Turrieta for assistance with experiments. This work was supported by NIH grants R35 GM131714 to Y.Z. and 5R01NS111749, 5R01DK109204, and 5R01EY10291 to H.E.H.

## AUTHOR CONTRIBUTIONS

Conceptualization: A.K., J.Y., H.E.H., and Y.Z.; Methodology: A.K., J.Y., and Y.Z.; Investigation: A.K., J.Y., L.A., Z.X., and L.L.; Resources: A.K., Y.Z., A.R.E., and H.E.H.; Data curation: A.K. and Y.Z.; Formal analysis: A.K. and Y.Z.; Software: A.K. and Y.Z.; Writing - original draft: A.K. and Y.Z.; Writing - review and editing: A.K., L.A., A.R.E., Z.X., L. L., H.E.H., and Y.Z.; Visualization: A.K. and Y.Z.; Supervision: Y.Z.; Project administration: Y.Z.; Funding acquisition: Y.Z. and H.E.H.

## DECLARATION OF INTERESTS

The authors declare no competing interests.

Declaration of generative AI use: ChatGPT was used to improve language during manuscript preparation and to assist with code development for data analysis and simulations. The authors reviewed and edited all AI-assisted text and code and take full responsibility for the content of this publication.

Data, code, and materials availability: All data needed to evaluate the conclusions are provided in the paper and/or Supplementary Materials. This study generated no new unique materials. Custom analysis and simulation code will be made available before publication.

## SUPPLEMENTAL INFORMATION

This PDF file contains Supplementary Text, Figures S1 to S20, Table S1, and References.

## METHOD DETAILS

### Protein sequences and purification

The amino acid sequences of the primary SNARE constructs are listed and annotated in Supplementary Text, Section T1, and Table S1. Genes encoding the cytosolic domains of syntaxin-1A and VAMP2 were cloned into pET-SUMO and pET-15b, respectively. Constructs containing the transmembrane domains of syntaxin-1A and VAMP2 were subcloned into pET-15b and pET-28a(+), respectively. SNAP-25 carrying a noncleavable N-terminal His tag was cloned into pET-15b. Recombinant proteins were expressed in Escherichia coli BL21-Gold (DE3) and purified by immobilized metal affinity chromatography on Ni-NTA agarose. Bound proteins were eluted with 300 mM imidazole and exchanged into storage buffer containing 25 mM HEPES (pH 7.4), 140 mM KCl, 2% (w/v) n-octyl-β-D-glucopyranoside (OG), and 2 mM tris(2-carboxyethyl)phosphine (TCEP). OG was omitted for protein constructs lacking transmembrane domains. For constructs with cleavable affinity tags, TEV-or SUMO-linked His tags were removed by incubation with the corresponding protease at a protein-to-protease molar ratio of 50:1. Released tags and proteases were removed by passage over fresh Ni-NTA resin, and tag-free proteins were collected in the flow-through.

Recombinant Gβ_1_γ_2_ was expressed in Sf9 insect cells using a baculovirus expression system as described previously^31^, with minor modifications. Forty-eight hours after infection, cells were harvested, lysed, and membrane fractions were isolated by centrifugation. Membrane proteins were solubilized with sodium cholate and purified by Ni-NTA affinity chromatography, followed by Mono Q anion-exchange chromatography. Purified Gβ1γ2 was concentrated, aliquoted, flash-frozen, and stored at −80 °C in CHAPS-containing buffer. C-terminally His-tagged Gα was purified using a protocol similar to that described for SNARE proteins. Purified nucleotide-free Gα was loaded with either GDP or the non-hydrolyzable GTP analog GTPγS prior to single-molecule experiments. For GDP loading, nucleotide-free Gα was incubated with 100 µM GDP for 3–4 h at 4 °C in the presence of 2 mM MgCl₂. For GTPγS loading, nucleotide-free Gα was incubated directly with 100 µM GTPγS for 3–4 h at 4 °C, after which MgCl₂ was added to a final concentration of 2 mM to stabilize the nucleotide-bound state. For optical tweezers experiments, the SNARE complex and Gβγ were co-reconstituted into SUVs using the standard reconstitution method and purified by density-gradient flotation, as described above. The GDP-or GTPγS-loaded Gα proteins were subsequently introduced into the microfluidic chamber at a final concentration of 2.5 µM to investigate nucleotide-dependent regulation of Gβγ interactions with the SNARE complex.

### Biotinylation of syntaxin and SpyTag003 conjugation of VAMP2

All syntaxin-1A variants containing a C-terminal AviTag were enzymatically biotinylated with BirA as described previously^56^. Syntaxin-1A (40 μM; 420 μL total volume) was supplemented with 10× Biomix-A, 10× Biomix-B, and 10× D-biotin according to the manufacturer’s instructions (BirA500, Avidity), together with His-tagged BirA at a syntaxin-to-BirA molar ratio of 130:1. The reaction was incubated overnight at 4 °C with gentle rotation. BirA was removed by Ni-NTA adsorption, and the sample was dialyzed to remove free biotin. Biotinylation was verified by streptavidin-binding analysis using SDS-PAGE.

A polypeptide-oligonucleotide conjugate (SpyTag-Oligo) was used to link VAMP2 to a DNA handle (Figure 1B). The polypeptide <u>RGVPHIVMVDAYKRYK</u>GGSDGSGNGSNGGG, containing the underlined SpyTag003 sequence^33^, and the oligonucleotide 5’-GAGGGCGTACAGTTGTATGTACGTTGGCGAGTTT-3’ were chemically synthesized and conjugated through click chemistry between the polypeptide C terminus and the oligonucleotide 3’ end (GenScript, NJ). The conjugated SpyTag-Oligo was purified to >90%. Purified SNARE complexes were incubated overnight at 4 °C with a fivefold molar excess of SpyTag-Oligo to enable covalent coupling to the C-terminal SpyCatcher003 fused to VAMP2 through the SpyTag003/SpyCatcher003 system.

### Cis-SNARE complex assembly

The ternary SNARE complex was assembled by mixing syntaxin-1A, SNAP-25, and VAMP2 at a molar ratio of 0.8:1:0.7 in the presence of 2 mM TCEP. The mixture was incubated overnight at 4 °C with gentle rotation to promote formation of the parallel four-helix bundle. Assembled complexes were purified through the N-terminal 6×His tag on SNAP-25 by Ni-NTA affinity chromatography. Complexes containing mutant proteins were assembled and purified by the same procedure.

### Lipids

The following lipids were purchased from Avanti Research (AL): 1-palmitoyl-2-oleoyl-sn-glycero-3-phosphocholine (POPC; 16:0–18:1 PC), 1,2-dioleoyl-sn-glycero-3-phospho-L-serine (DOPS; 18:1 PS), 1,2-dioleoyl-sn-glycero-3-phospho-(1′-myo-inositol-4′,5′-bisphosphate) [18:1 PI(4,5)P_2_], 1,2-dimyristoyl-sn-glycero-3-phosphocholine (DMPC; 14:0 PC), 1-stearoyl-2-arachidonoyl-sn-glycero-3-phosphocholine (SAPC; 18:0–20:4 PC), 1,2-distearoyl-sn-glycero-3-phosphoethanolamine-N-[biotinyl(polyethylene glycol)-2000] [DSPE-PEG(2000)-biotin; 18:0], 1,2-dioleoyl-sn-glycero-3-[(N-(5-amino-1-carboxypentyl)iminodiacetic acid)succinyl] nickel salt [DGS-NTA (Ni); 18:1], 1-palmitoyl-2-[12-[(7-nitro-2-1,3-benzoxadiazol-4-yl)amino]dodecanoyl]-sn-glycero-3-phosphocholine (NBD-PC; 16:0-12:0) and cholesterol. Lipids dissolved in chloroform were stored at −20 °C and used without further purification.

### Reconstitution of cis-SNAREs into SUVs and LUVs

For cis-SNARE pulling experiments, approximately 2 µmol of POPC was dried under nitrogen in a round-bottom glass vial and further desiccated under vacuum for at least 2 h. SNARE complexes were reconstituted into SUVs and LUVs by standard and direct methods, respectively, as described previously^39,40^, with minor modifications. For the standard method, the dried lipid film was rehydrated with 400 μL of buffer A [25 mM HEPES, 140 mM KCl, and 2% (w/v) OG] containing approximately 35 pmol of preassembled SNARE complex. The lipid–detergent–protein mixture was mixed vigorously for 30 min at room temperature. Detergent-free buffer B (600 μL; 25 mM HEPES and 140 mM KCl) was then added to a final volume of 1 mL. The sample was dialyzed overnight against 1 L of buffer B, with two buffer exchanges after 6 h, to remove detergent and promote vesicle formation. The dialyzed sample was purified by Nycodenz density-gradient flotation to isolate proteoliposomes containing reconstituted SNARE complexes.

In the direct method, dried lipid films were rehydrated in 1 mL of buffer B and vortexed for 30 min to form multilamellar vesicles. The suspension was subjected to at least seven freeze–thaw cycles, and then passed 21 times through a 100-nm-pore polycarbonate membrane (Cytiva, DE) using a mini-extruder (Avanti Research, AL) to generate LUVs. Vesicles were destabilized with Triton X-100 at a final concentration of 0.1% (w/v) and mixed end-over-end for 2 h at room temperature. SNARE complexes in buffer A were added and mixed for an additional 2 h under the same conditions. The detergent concentration was maintained at or below 0.1% (w/v) during protein incorporation. Detergent was then removed overnight at 4 °C with Bio-Beads SM-2 (Bio-Rad, CA) under gentle mixing, using a detergent-to-bead ratio of ∼0.1% (w/w). SUV and LUV proteoliposomes were purified by Nycodenz density-gradient flotation to isolate vesicles containing reconstituted SNARE complexes.

### Dithionite quenching assay

Proteoliposomes were prepared from 99.5 mol% POPC and 0.5 mol% NBD-PC and reconstituted with SNARE complexes as described above. NBD fluorescence was measured with a FluoroMax Plus spectrofluorometer (Horiba, Ltd.). After recording the initial fluorescence, freshly prepared sodium dithionite was added to 5 mM, decreasing fluorescence by approximately 50% (Figure S2A), consistent with selective quenching of NBD fluorophores in the outer leaflet. To probe the inner leaflet, 1% Triton X-100 and an additional 5 mM dithionite were added to solubilize the vesicles and completely quench the remaining fluorescence.

### Vesicle size measurements from optical tweezers and dynamic light scattering

To determine vesicle size after each cis-SNARE pulling experiment (Figure S4), SNARE complexes were assembled with syntaxin-1A and VAMP2 constructs lacking the engineered −6-layer cysteine substitutions, together with SNAP-25. Vesicle sizes were also measured by dynamic light scattering with a DynaPro NanoStar II instrument (Wyatt Technology, CA).

### Chymotrypsin cleavage assay and SNARE orientation on SUVs/LUVs

The orientation of SNARE complexes reconstituted into SUVs and LUVs was assessed by chymotrypsin proteolysis. Proteoliposomes were incubated with chymotrypsin at a 5:1 molar ratio relative to total SNARE complex for 30 min at room temperature. Chymotrypsin selectively cleaves protein regions exposed on the exterior of intact vesicles, whereas lumen-facing regions remain protected. Reactions were terminated with 1 mM phenylmethylsulfonyl fluoride (PMSF). Two controls were performed in parallel. For the negative control, 1 mM PMSF was added before chymotrypsin to inhibit proteolysis. For the positive control, proteoliposomes were solubilized with 2% OG before chymotrypsin treatment to expose all protein domains. After proteolysis, SDS-PAGE loading buffer was added, and samples were immediately heated at 95 °C for 5 min. Equal volumes of untreated, chymotrypsin-treated, negative control, and positive control samples were analyzed by SDS-PAGE. Band intensities were quantified with ImageJ (Figure S2).

### Preparation of trans-SNARE complexes

A protocol for preparing trans-SNARE complexes is shown in Figure S15. The t-SNARE complex was assembled by mixing syntaxin-1A, SNAP-25, and the C-terminal VAMP2 fragment (residues 49 to 96; Vc peptide) at a molar ratio of 1:1.2:3 in buffer A and incubating the mixture overnight at 4 °C. The stabilized t-SNARE acceptor complex was purified through the His tag on SNAP-25 by Ni-NTA affinity chromatography. For SUV reconstitution, the purified t-SNARE complex was mixed with 99.5 mol% POPC and 0.5 mol% DSPE-PEG(2000)-biotin in buffer A and dialyzed overnight against 1 L of detergent-free buffer B, with two buffer exchanges after 6 h, to remove detergent and drive proteoliposome formation. In parallel, VAMP2-containing nanodiscs were assembled by mixing membrane scaffold protein MSP1E3D1 (mean diameter, ∼13 nm), 1 µM POPC, and VAMP2 at a molar ratio of 2:120:0.05 in buffer A. Detergent was removed by overnight incubation with Bio-Beads at 4 °C under gentle agitation [detergent-to-bead ratio, 0.1% (w/w)]. Nanodiscs were purified through the His tag on VAMP2 to remove empty nanodiscs. To generate trans-SNARE complexes, VAMP2-containing nanodiscs were mixed with t-SNARE-reconstituted SUVs at an approximately 1:1 molar ratio in buffer B and incubated at room temperature or 4 °C. Full-length VAMP2 displaced the Vc peptide and formed a trans-SNARE complex bridging the nanodisc and SUV membranes. The assemblies were purified by Nycodenz density-gradient flotation to remove free VAMP2 and aggregates.

### High-resolution dual-trap optical tweezers

Dual-trap optical tweezers were built as described previously^57,58^. A 1064-nm laser was expanded, collimated, and divided into two orthogonally polarized beams. One beam was reflected from a mirror mounted on a nanopositioning stage (Mad City Labs, WI) to control trap position. The beams were recombined, expanded, and focused through a 60× water-immersion objective (NA 1.2; Olympus, PA) to form two traps in the central channel of a custom microfluidic flow chamber. One trap was stationary, and the other was moved by the nanopositioning stage. Light from the traps was collected by a second water-immersion objective, separated by polarization, and detected by two position-sensitive detectors (Pacific Silicon Sensor, CA). Bead displacement was measured by back-focal-plane interferometry, and trap stiffness was calibrated from the Brownian motion of trapped beads. The instrument was controlled through a custom LabVIEW interface (National Instruments, TX).

### Single-molecule experiment

A detailed protocol for single-molecule optical-tweezers experiments is shown in Figure S1 and has been described elsewhere^56^. An aliquot of proteoliposomes was incubated sequentially with DNA handle 1 and 5 μL of anti-digoxigenin-coated polystyrene beads (2.17 μm in diameter; Spherotech, IL). DNA handle 2 was incubated with tetrameric streptavidin (Promega, WI) and then with 5 μL of anti-digoxigenin-coated polystyrene beads. Each bead mixture was diluted in 1 mL of buffer B and injected separately into the top or bottom channel of the microfluidic chamber. For cytosolic SNARE pulling experiments, streptavidin-coated beads (1.86 μm in diameter) tethered the complex through biotinylated syntaxin-1A. The top and bottom channels were connected by capillaries to a central channel, where both bead types were trapped. A single SNARE complex was tethered between two beads by bringing the beads into proximity. Data were acquired at 20 kHz, mean-filtered to 10 kHz, and stored for analysis. Experiments were performed in buffer B at 23 ± 1 °C. An oxygen-scavenging system was included to minimize photodamage from the optical traps^56^. Force was applied to single tethers by increasing or decreasing the trap separation at 10 nm s^-1^.

### Data analysis

Detailed procedures for analyzing single-molecule trajectories are described elsewhere^12,36,56^. Extension trajectories acquired at constant trap separations or mean forces were analyzed by hidden Markov modeling to determine state probabilities, forces, extensions, transition rates, and idealized extension trajectories (Figures S13 and S20)^35^. The three states were ordered sequentially by extension, with transition rates between nonadjacent states typically orders of magnitude lower than those between adjacent states. These sequential transitions support the one-dimensional energy landscape used to describe SNARE zippering. These measurements were fitted with a force-induced protein folding model to obtain a simplified energy landscape at zero force or under load^36^ (Figure 1G and Figure S14). The model used the zero-force energy landscape together with DNA handle lengths, polypeptide linkers, and trap stiffnesses. The landscape was parameterized by energy minima representing folding states and maxima representing transition states, each associated with a contour length of unfolded polypeptide. The three-state SNARE transition, for example, was described by six free parameters after assigning the known contour length and zero energy to the unzipped state. State contour lengths were assumed to be force-independent. The model predicted state probabilities and transition rates at each trap separation or mean force using the Boltzmann distribution and Kramers’ theory, respectively^36^. The measured extension of the protein–DNA tether was modeled as the sum of the extensions of the DNA handles and the SNARE complex along the pulling direction; the latter comprises structured and unfolded portions. Extensions of unfolded polypeptide and DNA handles were calculated with worm-like-chain models for semiflexible polymers. Extensions of structured SNARE portions were derived from the crystal structure of the fully assembled complex and the geometry of TMD tilting (Figure S12 and Supplementary Text), assuming the folding and unfolding pathways defined in Figure 1E and Figure S7A. Specifically, VAMP2 sequentially zippers and unzips on a structured t-SNARE template between the crosslinking site at T35 and the LD C-terminus at M95 (Figure 1G, inset). The t-SNARE template was assumed to adopt the structure observed in the fully assembled SNARE complex^13^. Energy-landscape parameters were obtained by nonlinear least-squares fitting of model predictions to the measurements. Parameters from 1 to 13 force scans were averaged; Figure S13 shows an example fit from one scan. Unless otherwise indicated, values are reported as mean ± SEM.

Unless otherwise indicated, the extension and force data shown in all FECs and extension– time trajectories were filtered using a 10-ms moving-average window.

### Trans-SNARE complex simulations

The trans-SNARE energy function followed the protein-folding model described above and is detailed in Supplementary Text, Sections T4 and T5.

### Quantification and statistical analysis

Data are shown as mean±S.D. or mean±SEM. For optical-tweezers experiments, repeated pulling cycles, force scans, and transition or dwell events from the same tether were treated as repeated measurements and averaged before group-level analysis. Constant-force trajectories were analyzed by hidden Markov modeling, and energy landscapes were obtained by nonlinear least-squares fitting to a force-dependent folding model. Gβγ-clamped-state survival probabilities were fitted with single- or biexponential functions, using a minimum dwell-time threshold of 0.5 s for manual identification. No statistical method was used to predetermine sample size, and no numerical outliers were excluded. Sample sizes were based on previous single-molecule force-spectroscopy studies and the number of measurements required to resolve conformational states and transition kinetics. All numerical analyses, model fitting, and simulations were performed in MATLAB.

## Notes

### Competing Interest Statement

The authors have declared no competing interest.

## REFERENCES

1. Alabi, A.A. & Tsien, R.W. Perspectives on kiss-and-run: role in exocytosis, endocytosis, and neurotransmission. Annu. Rev. Physiol. 75, 393–422 (2013).

2. Jahn, R., Cafiso, D.C. & Tamm, L.K. Mechanisms of SNARE proteins in membrane fusion. Nat. Rev. Mol. Cell Biol. 25, 101–118 (2023).

3. Tao, C.L. et al. “Kiss-shrink-run” unifies mechanisms for synaptic vesicle exocytosis and hyperfast recycling. Science 390, eads7954 (2025).

4. Sudhof, T.C. & Rothman, J.E. Membrane fusion: grappling with SNARE and SM proteins. Science 323, 474–477 (2009).

5. Rizo, J. Molecular mechanisms underlying neurotransmitter release. Annu. Rev. Biophys. 51, 377–408 (2022).

6. Jahn, R. & Scheller, R.H. SNAREs - engines for membrane fusion. Nat. Rev. Mol. Cell Bio. 7, 631–643 (2006).

7. Blackmer, T., Larsen, E.C., Takahashi, M., Martin, T.F.J., Alford, S. & Hamm, H.E. G protein âã subunit-mediated presynaptic inhibition: Regulation of exocytotic fusion downstream of Ca^2+^ entry. Science 292, 293–297 (2001).

8. Zhang, Y.L. & Hughson, F.M. Chaperoning SNARE folding and assembly. Annu. Rev. Biochem. 90, 581–603 (2021).

9. Yim, Y.Y., Zurawski, Z. & Hamm, H. GPCR regulation of secretion. Pharmacol. Ther. 192, 124–140 (2018).

10. Hanson, P.I., Roth, R., Morisaki, H., Jahn, R. & Heuser, J.E. Structure and conformational changes in NSF and its membrane receptor complexes visualized by quick-freeze/deep-etch electron microscopy. Cell 90, 523–535 (1997).

11. Sutton, R.B., Fasshauer, D., Jahn, R. & Brunger, A.T. Crystal structure of a SNARE complex involved in synaptic exocytosis at 2.4 angstrom resolution. Nature 395, 347–353 (1998).

12. Gao, Y., Zorman, S., Gundersen, G., Xi, Z.Q., Ma, L., Sirinakis, G., Rothman, J.E. & Zhang, Y.L. Single reconstituted neuronal SNARE complexes zipper in three distinct stages. Science 337, 1340–1343 (2012).

13. Stein, A., Weber, G., Wahl, M.C. & Jahn, R. Helical extension of the neuronal SNARE complex into the membrane. Nature 460, 525–528 (2009).

14. Min, D., Kim, K., Hyeon, C., Cho, Y.H., Shin, Y.K. & Yoon, T.Y. Mechanical unzipping and rezipping of a single SNARE complex reveals hysteresis as a force-generating mechanism. Nat. Commun. 4, 1705 (2013).

15. Lai, Y. et al. Molecular mechanisms of synaptic vesicle priming by Munc13 and Munc18. Neuron 95, 591–607 (2017).

16. Kiessling, V., Kreutzberger, A.B., Liang, B.Y., Nyenhuis, S.B., Seelheim, P., Castle, J.D., Cafiso, D.S. & Tamm, L.K. A molecular mechanism for calcium-mediated synaptotagmin-triggered exocytosis. Nat. Struct. Mol. Biol. 25, 911–917 (2018).

17. Han, X., Wang, C.T., Bai, J.H., Chapman, E.R. & Jackson, M.B. Transmembrane segments of syntaxin line the fusion pore of Ca^2+^-triggered exocytosis. Science 304, 289–292 (2004).

18. Sharma, S. & Lindau, M. Molecular mechanism of fusion pore formation driven by the neuronal SNARE complex. Proc. Natl. Acad. Sci. USA 115, 12751–12756 (2018).

19. Rathore, S.S., Liu, Y.H., Yu, H.J., Wan, C., Lee, M., Yin, Q., Stowell, M.H.B. & Shen, J.S. Intracellular vesicle fusion requires a membrane-destabilizing peptide located at the juxtamembrane region of the v-SNARE. Cell Rep. 29, 4583–4592.e3 (2019).

20. Rizo, J., Chattopadhyay, M., Wosztyl, A. & Xu, J.J. The local detergent model of SNARE-mediated membrane fusion. J. Cell Sci. 139, jcs264344 (2026).

21. Dhara, M. et al. v-SNARE transmembrane domains function as catalysts for vesicle fusion. Elife 5, e17571 (2016).

22. Zhang, Y., Hu, Y., Xie, H., Liu, M., Yang, J. & Ma, C. Structural mechanism of soluble N-ethylmaleimide-sensitive factor attachment protein receptor complex assembly in lipid bilayers revealed by solid-state NMR. J. Am. Chem. Soc. 145, 10641–10650 (2023).

23. Weber, T., Zemelman, B.V., McNew, J.A., Westermann, B., Gmachl, M., Parlati, F., Sollner, T.H. & Rothman, J.E. SNAREpins: Minimal machinery for membrane fusion. Cell 92, 759–772 (1998).

24. Knecht, V. & Grubmuller, H. Mechanical coupling via the membrane fusion SNARE protein syntaxin 1A: A molecular dynamics study. Biophys. J. 84, 1527–1547 (2003).

25. Ma, L., Rebane, A.A., Yang, G., Xi, Z., Kang, Y., Gao, Y. & Zhang, Y.L. Munc18-1-regulated stage-wise SNARE assembly underlying synaptic exocytosis. Elife 4, e09580 (2015).

26. Li, F., Pincet, F., Perez, E., Eng, W.S., Melia, T.J., Rothman, J.E. & Tareste, D. Energetics and dynamics of SNAREpin folding across lipid bilayers. Nat. Struct. Mol. Biol. 14, 890–896 (2007).

27. Chernomordik, L.V. & Kozlov, M.M. Mechanics of membrane fusion. Nat. Struct. Mol. Biol. 15, 675–683 (2008).

28. Rizo, J., Sari, L., Jaczynska, K., Rosenmund, C. & Lin, M.M. Molecular mechanism underlying SNARE-mediated membrane fusion enlightened by all-atom molecular dynamics simulations. Proc. Natl. Acad. Sci. USA 121, e2321447121 (2024).

29. Mostafavi, H., Thiyagarajan, S., Stratton, B.S., Karatekin, E., Warner, J.M., Rothman, J.E. & O’Shaughnessy, B. Entropic forces drive self-organization and membrane fusion by SNARE proteins. Proc. Natl. Acad. Sci. USA 114, 5455–5460 (2017).

30. Zorman, S., Rebane, A.A., Ma, L., Yang, G.C., Molski, M.A., Coleman, J., Pincet, F., Rothman, J.E. & Zhang, Y.L. Common intermediates and kinetics, but different energetics, in the assembly of SNARE proteins. Elife 3, e03348 (2014).

31. Eitel, A.R. et al. Molecular basis for Gbetagamma-SNARE-mediated inhibition of synaptic vesicle fusion. J. Biol. Chem. 301, 110377 (2025).

32. Bustamante, C.J., Chemla, Y.R., Liu, S.X. & Wang, M.D. Optical tweezers in single-molecule biophysics. Nat. Rev. Methods Primers 1 25 (2021).

33. Keeble, A.H., Turkki, P., Stokes, S., Anuar, I.N.A.K., Rahikainen, R., Hytonen, V.P. & Howarth, M. Approaching infinite affinity through engineering of peptide-protein interaction. Proc. Natl. Acad. Sci. USA 116, 26523–26533 (2019).

34. Choi, H.K. et al. Watching helical membrane proteins fold reveals a common N-to-C-terminal folding pathway. Science 366, 1150–1156 (2019).

35. Zhang, Y.L., Jiao, J. & Rebane, A.A. Hidden Markov modeling with detailed balance and its application to single protein folding. Biophys. J. 111, 2110–2124 (2016).

36. Rebane, A.A., Ma, L. & Zhang, Y.L. Structure-based derivation of protein folding intermediates and energies from optical tweezers. Biophys. J. 110, 441–454 (2016).

37. Yamaga, M. & Martin, T.F.J. PI(4,5)P_2_ is a master regulator for Ca^2+^-triggered vesicle exocytosis. Biochim. Biophys. Acta 1870, 159651 (2025).

38. Rituper, B., Flasker, A., Gucek, A., Chowdhury, H.H. & Zorec, R. Cholesterol and regulated exocytosis: A requirement for unitary exocytotic events. Cell Calcium 52, 250–258 (2012).

39. Hernandez, J.M., Stein, A., Behrmann, E., Riedel, D., Cypionka, A., Farsi, Z., Walla, P.J., Raunser, S. & Jahn, R. Membrane fusion intermediates via directional and full assembly of the SNARE complex. Science 336, 1581–1584 (2012).

40. Shi, L., Shen, Q.T., Kiel, A., Wang, J., Wang, H.W., Melia, T.J., Rothman, J.E. & Pincet, F. SNARE proteins: One to fuse and three to keep the nascent fusion pore open. Science 335, 1355–1359 (2012).

41. Bao, H. et al. Dynamics and number of trans-SNARE complexes determine nascent fusion pore properties. Nature 554, 260–263 (2018).

42. Diao, J. et al. Synaptic proteins promote calcium-triggered fast transition from point contact to full fusion. Elife 1, e00109 (2012).

43. Vardar, G., Salazar-Lazaro, A., Zobel, S., Trimbuch, T. & Rosenmund, C. Syntaxin-1A modulates vesicle fusion in mammalian neurons via juxtamembrane domain dependent palmitoylation of its transmembrane domain. Elife 11, e78182 (2022).

44. McNew, J.A., Weber, T., Engelman, D.M., Sollner, T.H. & Rothman, J.E. The length of the flexible SNAREpin juxtamembrane region is a critical determinant of SNARE-dependent fusion. Mol. Cell 4, 415–421 (1999).

45. Wickner, W., Lopes, K., Song, H., Rizo, J. & Orr, A. Efficient fusion requires a membrane anchor on the vacuolar Qa-SNARE. Mol. Biol. Cell 34, ar88 (2023).

46. Kesavan, J., Borisovska, M. & Bruns, D. v-SNARE actions during Ca^2+^-triggered exocytosis. Cell 131, 351–363 (2007).

47. Zhou, P., Bacaj, T., Yang, X., Pang, Z.P. & Sudhof, T.C. Lipid-anchored snares lacking transmembrane regions fully support membrane fusion during neurotransmitter release. Neuron 80, 470–83 (2013).

48. D’Agostino, M., Risselada, H.J., Lurick, A., Ungermann, C. & Mayer, A. A tethering complex drives the terminal stage of SNARE-dependent membrane fusion. Nature 551, 634–638 (2017).

49. van den Bogaart, G., Holt, M.G., Bunt, G., Riedel, D., Wouters, F.S. & Jahn, R. One SNARE complex is sufficient for membrane fusion. Nat. Struct. Mol. Biol. 17, 358–364 (2010).

50. Shin, J., Lou, X.C., Kweon, D.H. & Shin, Y.K. Multiple conformations of a single SNAREpin between two nanodisc membranes reveal diverse pre-fusion states. Biochem. J. 459, 95–102 (2014).

51. Rost, B.R., Nicholson, P., Ahnert-Hilger, G., Rummel, A., Rosenmund, C., Breustedt, J. & Schmitz, D. Activation of metabotropic GABA receptors increases the energy barrier for vesicle fusion. J. Cell Sci. 124, 3066–3073 (2011).

52. Walter, A.M., Wiederhold, K., Bruns, D., Fasshauer, D. & Sorensen, J.B. Synaptobrevin N-terminally bound to syntaxin-SNAP-25 defines the primed vesicle state in regulated exocytosis. J. Cell Biol. 188, 401–413 (2010).

53. Yu, H., Shen, C., Liu, Y., Menasche, B.L., Ouyang, Y., Stowell, M.H.B. & Shen, J. SNARE zippering requires activation by SNARE-like peptides in Sec1/Munc18 proteins. Proc. Natl. Acad. Sci. USA 115, E8421–E8429 (2018).

54. Rebane, A.A., Wang, B., Ma, L., Qu, H., Coleman, J., Krishnakumar, S.S., Rothman, J.E. & Zhang, Y.L. Two disease-causing SNAP-25B mutations selectively impair SNARE C-terminal assembly. J. Mol. Biol. 430, 479–490 (2018).

55. Guzman, R.E., Schwarz, Y.N., Rettig, J. & Bruns, D. SNARE force synchronizes synaptic vesicle fusion and controls the kinetics of quantal synaptic transmission. J. Neurosci. 30, 10272–10281 (2010).

56. Jiao, J.Y., Rebane, A.A., Ma, L. & Zhang, Y.L. Single-molecule protein folding experiments using high-resolution optical tweezers. Methods Mol. Biol. 1486, 357–390 (2017).

57. Sirinakis, G., Ren, Y.X., Gao, Y., Xi, Z.Q. & Zhang, Y.L. Combined and versatile high-resolution optical tweezers and single-molecule fluorescence microscopy. Rev. Sci. Instrum. 83, 093708 (2012).

58. Moffitt, J.R., Chemla, Y.R., Izhaky, D. & Bustamante, C. Differential detection of dual traps improves the spatial resolution of optical tweezers. Proc. Natl. Acad. Sci. USA 103, 9006–9011 (2006).

