## Supplementary Information for "Membrane-gated SNARE zippering focuses energy for fusion"

This PDF file includes:

Supplementary Text
Figures S1 to S20
Table S1
References

### Supplementary Text

#### T1. Amino acid sequences of primary protein constructs

**Syntaxin-1A (L205C, C272S, C273S)**

MHHHHHHENLYFQS[1]*M*KDRTQELRTAKDSDDDDDVTVTDRDRFMDEFFEQVEEI
RGFIDKIAENVEEVKRKHSAILASPNPDEKTKEELEELMSDIKKTANKVRSKLKSIEQ
SIEQEEGLNRSSADLRIRKTQHSTLSRKFVEVMSEYNATQSDYRERSKGRIQRQLEIT
GRTTTSEELEDMLESGNPAIFAS [180]*G* IIMDSSISKQALSEIETRHSEIIK [205]***C*** ENSIRELHD
MFMDMAMLVESQGEMIDRIEYNVEHAVDYVERAVSDTKKAV [256]*K* YQSKARRK [265]*K* I
MIII**SS**VILGIIIASTIGGIF [288]*G* GGSGNGRGGSDEGSQGDNGSGDGSKGSGNESGQGTG
EGSNGSGDGSGQGTGHSGNGSGQENGDQGSGRGGSGSKGGSGGGSGGLNDIFEAQ
KIEWHE

Black underlining and cyan highlighting indicate the tobacco etch virus (TEV) protease cleavage site and AviTag, respectively. The Habc domain and SNARE motif of syntaxin-1A are highlighted in black, whereas the linker domain (LD) and transmembrane domain (TMD) are shown in green and magenta, respectively. The two native cysteines at positions 272 and 273 in the TMD were replaced by the bold serine residues to facilitate tether formation. An 83-residue flexible linker (gray) was added after the TMD to facilitate membrane-based pulling experiments.

**Cytosolic syntaxin-1A (L205C, no TMD)**

MGSSHHHHHHSSGLVPRGSHMASMSDSEVNQEAKPEVKPEVKPETHINLKVSDGSS
EIFFKIKKTTPLRRLMEAFAKRQGKEMDSLRFLYDGIRIQADQTPEDLDMEDNDIIEA
HREQIGGS[1]*M*KDRTQELRTAKDSDDDDDVTVTDRDRFMDEFFEQVEEIRGFIDKIA
ENVEEVKRKHSAILASPNPDEKTKEELEELMSDIKKTANKVRSKLKSIEQSIEQEEGL
NRSSADLRIRKTQHSTLSRKFVEVMSEYNATQSDYRERSKGRIQRQLEITGRTTTSEE
LEDMLESGNPAIFAS [180]*G* IIMDSSISKQALSEIETRHSEIIK [205]***C*** ENSIRELHDMFMDMAM
LVESQGEMIDRIEYNVEHAVDYVERAVSDTKKAV [256]*K* YQSKARRK [265]*K* GGSGNGGSG
SGLNDIFEAQKIEWHE

Black underlining and yellow and cyan highlighting denote the thrombin cleavage site, SUMO tag, and AviTag, respectively. The Habc domain and SNARE motif of syntaxin-1A are highlighted in black, and the LD is highlighted in green. Lys205 at the −6 layer was mutated to cysteine to enable site-specific crosslinking with VAMP2. A 10-residue flexible linker (gray) was introduced immediately after the LD to facilitate attachment of the DNA handle to syntaxin.

**VAMP2 (Q36C, C103S)**

MGSSHHHHHHGSGLVPRGSASMSDSEVNQEAKPEVKPEVKPETHINLKVSDGSSEI
FFKIKKTTPLRRLMEAFAKRQGKEMDSLRFLYDGIRIQADQTPEDLDMEDNDIIEAH

1 36
REQIGGSS*M*SATAATVPPAAPAGEGGPPAPPNLTSNRRLQQT***C***AQVDEVVDIMRV
85 94
NVDKVLERDQKLSELDDRADALQAGASQFETSAAKL*K*RKYWWKNL*K*MMIILGVI
116
SAIILIIIIVYFS *T* GSGGRGNGGSAGSKGGSGGHGVTTLSGLSGEQGPSGDMTTEEDS
ATHIKFSKRDEDGRELAGATMELRDSSGKTISTWISDGHVKDFYLYPGKYTFVETA
APDGYEVATPIEFTVNEDGQVTVDGEATEGDAHTG

Black underlining and yellow and orange highlighting denote the thrombin cleavage site, SUMO tag, and SpyCatcher003, respectively. The SNARE motif, LD, and TMD of VAMP2 are highlighted in black, green, and magenta, respectively. A 21-residue flexible linker (gray) was introduced immediately after the TMD to facilitate pulling.

**Cytosolic VAMP2 (no TMD)**
MGSSHHHHHHGSG<u>LVPRGS</u>ASMSDSEVNQEAKPEVKPEVKPETHINLKVSDGSSEI
FFKIKKTTPLRRLMEAFAKRQGKEMDSLRFLYDGIRIQADQTPEDLDMEDNDIIEAH
1 36
REQIGGSS*M*SATAATVPPAAPAGEGGPPAPPNLTSNRRLQQT***C***AQVDEVVDIMRV
85 94
NVDKVLERDQKLSELDDRADALQAGASQFETSAAKL*K*RKYWWKNL*K*MGSGGRG
NGGSAGSKGGSGGHGVTTLSGLSGEQGPSGDMTTEEDSATHIKFSKRDEDGRELAG
ATMELRDSSGKTISTWISDGHVKDFYLYPGKYTFVETAAPDGYEVATPIEFTVNEDG
QVTVDGEATEGDAHTG

Black underlining and yellow and orange highlighting denote the thrombin cleavage site, SUMO tag, and SpyCatcher003, respectively. The SNARE motif and LD of VAMP2 are highlighted in black and green, respectively. Gln36 at the −6 layer was replaced by cysteine to enable crosslinking with syntaxin. A 21-residue flexible linker (gray) was introduced immediately after the LD to facilitate pulling.

**SNAP-25B (C85S, C88S, C90S, C92S)**
MAHHHHHHMAEDADMRNELEEMQRRADQLADESLESTRRMLQLVEESKDAGIRT
LVMLDEQGEQLERIEEGMDQINKDMKEAEKNLTDLGKFSGLSVSPSNKLKSSDAYK
KAWGNNQDGVVASQPARVVDEREQMAISGGFIRRVTNDARENEMDENLEQVSGII
GNLRHMALDMGNEIDTQNRQIDRIMEKADSNKTRIDEANQRATKMLGSG

To prevent nonspecific crosslinking with syntaxin-1A or VAMP2, the four native cysteine residues in SNAP-25B at positions 85, 88, 90, and 92 were replaced with serine.

### T2. Preparation of DNA handles

Two 2,249-bp DNA handles were used in the optical tweezers experiments. Each handle contained two digoxigenin labels at one end and either a single-stranded overhang or biotin at the other. Both handles were generated by polymerase chain reaction (PCR) using the primers listed below.
<u>Handle 1:</u>
Forward primer: 5′-[2xDIG]TCGCCACCA[DIG]TCATTTCCAGCTTTTGTG-3′

Reverse primer: 5′-CTCG<u>CCAACGTACATACAACTGTACGCCCTC</u>-S18-ACTATCGCCACTTTTATTGGCG-3′

Handle 2:

Forward primer: 5′-[2xDIG]TCGCCACCA[DIG]TCA TTT CCA GCT TTT GTG-3′

Reverse primer: 5′-Biotin-ATCATCCAAGGCTGAGCCTGCAG-3′

S18 is an 18-atom hexaethylene glycol spacer that prevents polymerase extension into the overhang region (underlined) during PCR. Lambda DNA (New England Biolabs, MA) served as the template for both handles, and primers were obtained from Integrated DNA Technologies.

**T3. Extension contribution of transmembrane domains**

The extension of the SNARE complex measured between the C termini of syntaxin and VAMP2 $X$ equals the extension of the structured cytosolic SNARE complex inside the SUV $x_c$ plus the projections of the two transmembrane helices along the pulling direction $x_h$, or

$$X = x_c + x_h, \tag{1}$$

with

$$x_h = 2h\sin\left(\theta + \theta_h\right), \tag{2}$$

where $h$ is the length of each transmembrane helix and $\theta_h$ and $\theta + \theta_h$ are the corresponding tilt angles relative to the membrane normal and the vertical direction, respectively (Figure S12A with $\theta_h = 0$). Note that

$$\sin\theta = \frac{x_c}{2R}, \tag{3}$$

where $R$ is the radius of the SUV. In the fully assembled SNARE complex, the two transmembrane helices are approximately parallel to each other and perpendicular to the membrane[1]. Therefore, the extension change due to TMD tilting is

$$x_h = 2h\sin\left[\arcsin\left(\frac{x}{2R}\right) + \theta_h\right]. \tag{4}$$

The tilt angle $\theta_h$ is generally small under our experimental conditions, as estimated below. Assuming $\theta_h = 0$, Eq. (4) simplifies to

$$x_h = \frac{hx}{R}. \tag{5}$$

To estimate the extension, we used $R =$ 22 nm, corresponding to the average radius of SUVs used in our experiments, $h = 5$ nm, and the maximum extension change $x_c$ as 18 nm for the unzipped cis-SNARE complex, corresponding to the N-terminal crosslinking site at VAMP2 residue 35. Based on Eq. (5), the estimated maximum extension change due to TMD tilting is 4.1 nm. For the fully zippered SNARE complex, the TMD separation was set to zero, with the last unfolded VAMP2 residue 95. The TMD extension as a function of the intermediate unfolded VAMP2 residue number was calculated as the linear interpolation of the two boundary points, yielding

$$x_h = -0.0683 \times AA + 6.49, \quad (6)$$

where $AA$ is the residue number of the last unfolded VAMP2 residue, with $35 \le AA \le 95$. The total extension of the structured portion of the cis-SNARE complex is

$$H = H_s + x_h \quad (7)$$

where $H_s$ is the extension of the structured portion of the cytosolic SNARE complex calculated from the crystal structure of the fully assembled SNARE complex (Figure S12).

To justify the assumption that pulling only slightly tilts the transmembrane helices under our experimental conditions, we considered the total energy of a TMD being pulled from its two ends in a parallel direction (Figure S12):

$$E = \frac{1}{2}k\left(\theta_h\right)^2 - Fx_h = \frac{1}{2}k\left(\theta_h - \theta\right)^2 - Fh\sin\left(\theta + \theta_h\right), \quad (8)$$

where $k$ is the tilt force constant of the transmembrane helix, chosen as 70 cal $mol^{-1}$ $deg^{-2}$ $=1.6\times10^3$ pN×nm $rad^{-2}$ and $F$ is the pulling force.[2] The equilibrium helix tilt angle was determined by minimizing the energy with respect to $\theta_h$, yielding

$$\theta_h = \frac{Fh}{k}\cos\left(\theta_h + \theta\right). \quad (9)$$

Numerical calculations indicated that at a representative pulling force of 15 pN, the helix tilt away from the membrane normal is ~3°, substantially smaller than the maximum helix tilt angle $\theta$ ≈32° caused by SNARE unfolding. Force-induced TMD tilting away from the membrane normal is therefore negligible under our experimental conditions.

#### T4. Membrane repulsive force and energy

Surface force apparatus (SFA) measurements reported the repulsive force between two curved protein-free membranes as

$$\frac{F(d)}{R} \approx \left(20 - 30\ \mathrm{Nm^{-1}}\right)e^{-d/(0.8\ \mathrm{nm})}, \quad (1)$$

where $F(d)$ is the force as a function of membrane separation $d$ and $R = 2$ cm is the radius of curvature of the two mica surfaces coated with membranes.[3] An approximate fit to these data is

$$\frac{F(d)}{R} \approx 25e^{-d/(0.8\ \mathrm{nm})}\ \mathrm{mNm^{-1}}. \quad (2)$$

The interaction free energy per unit area $e$ between two equivalent planar membrane surfaces is

$$e = \frac{F}{2\pi R} \approx e^{-d/(0.8\ \mathrm{nm})}\ \mathrm{k_BT/nm^2}. \quad (3)$$

For SNARE-anchored membranes, the repulsive force decays with membrane separation $d$ with apparent decay lengths of ~8 nm at separations of 9 to 20 nm and ~2.5 nm at separations of 2 to 6 nm. A plateau separates the two regions associated with SNARE assembly. An approximate fit to the measured force data is

$$\frac{F(d)}{R} \approx 18e^{-d/8}\quad \left(9 - 20\ \mathrm{nm}\right) \quad (4)$$

and

$$\frac{F(d)}{R} \approx 55e^{-d/2.5} \quad (2-6 \text{ nm}), \tag{5}$$

with $d$ in nm and $F/R$ in pN×nm$^{-1}$. The corresponding interaction energy per unit area is approximately

$$e(d) \approx 0.71e^{-d/8} \quad (9-20 \text{ nm}) \tag{6}$$

and

$$e(d) \approx 2.15e^{-d/2.5} \quad (2-6 \text{ nm}) \tag{7}$$

in $k_B T$ /nm².

Whether biological membranes exhibit such long-range interactions is unclear. The long-range repulsive interactions between SNARE-containing membranes might arise from immobilized SNARE proteins on supported bilayers in the SFA measurements[3]. Such interactions may be absent when membrane proteins are freely mobile. Alternatively, the long-range repulsion may arise from trans-SNARE complexes projecting from the membrane[4]. In our simulation, we therefore used the decay length of 0.8 nm measured for protein-free membranes.

We calculated the total repulsive energy between a spherical vesicle of radius $R$ and a flat plasma membrane (PM) by integrating the interaction energy density over the projected contact area. If the closest separation between the vesicle and PM surfaces is $d$, the total energy is

$$E_m(d) = 2\pi \int_0^R e(z) r dr \tag{8}$$

where

$$z = d + R - \sqrt{R^2 - r^2} \tag{9}$$

and

$$e(z) = Ae^{-\frac{z}{d_m}} \tag{10}$$

with $A \approx 1 \text{ k}_\text{B}\text{T/nm}^2$ according to Eq. (3). Substitution of Eqs. (9) and (10) into Eq. (8) yields

$$E_m(d) = A_m e^{-\frac{d}{d_m}}, \tag{11}$$

where $A_m$ is the maximum repulsive energy, with

$$A_m = 2\pi A d_m \left[ d_m e^{-\frac{R}{d_m}} + (R - d_m) \right] \approx 2\pi A d_m R. \tag{12}$$

Therefore, the repulsive membrane energy increases approximately linearly with vesicle radius. Vesicles with radii of 20 nm or 50 nm, for example, have maximum repulsive energies $A_m$ of approximately 100 $k_BT$ or 250 $k_BT$, respectively. The repulsive force between the vesicle and the flat membrane is

$$F(d) = -\frac{dE_m}{dd} = \frac{A_m}{d_m} e^{-\frac{d}{d_m}}. \tag{13}$$

The maximum repulsive force at contact is approximately 513 pN for a 20-nm-radius vesicle and 1281 pN for a 50-nm-radius vesicle. These estimates assume that both the vesicle and PM remain undeformed during close apposition. The actual energy barrier during SNARE-mediated fusion may differ because local membrane deformation, lipid rearrangement, and protein organization can substantially alter the fusion pathway.

**T5. Conformations and energetics of trans-SNARE complexes**

Consider $N_s$ identical trans-SNARE complexes bridging a flat membrane representative of the plasma membrane and a vesicle of radius $R$ in an approximately circularly symmetric arrangement (Figure S16A illustrates one trans-SNARE complex). Because of repulsion between the PM and the vesicle membrane, the vesicle equilibrates at the closest distance $d$ from the PM. We modeled the partially zippered SNARE complex as a rod of length $L$, corresponding to the length of the structured t-SNARE complex, tilted relative to the PM by a fixed angle $\alpha$ (Figure S16E). Based on the crystal structure of the SNARE complex[1], the length $L$ was chosen as 12 nm, corresponding to the distance between Cα positions of syntaxin K265 and SNAP-25B R8. The C termini of syntaxin and VAMP2 are anchored in the PM at point A and the vesicle at point E, respectively, via their transmembrane domains. The trans-SNARE folding state is characterized by the contour length $l$ of VAMP2 unfolded or peeled from the SNARE rod from point A to point D, with the associated SNARE folding energy $V(l)$. The extension of the unfolded VAMP2 polypeptide $x_v = |DE|$ is related to the force exerted by the membrane $F$ and the contour length $l$, which can be described by the Marko–Siggia formula[5]

$$F = \frac{k_B T}{P}\left[\frac{1}{4\left(1-\frac{x_v}{l}\right)^2} + \frac{x_v}{l} - \frac{1}{4}\right], \tag{14}$$

where $k_B T = 4.1$ pN×nm and $P$ is the persistence length of the polypeptide, with $P = 0.6$ nm. The associated entropic energy of polypeptide stretching is

$$E_s\left(x_v, l\right) = \frac{k_B T}{P}\frac{l}{4\left(1-\frac{x_v}{l}\right)}\left[3\left(\frac{x_v}{l}\right)^2 - 2\left(\frac{x_v}{l}\right)^3\right]. \tag{15}$$

The associated length change $h = |AD|$ varies approximately linearly with the contour length, or

$$h = al + b, \tag{16}$$

where $a$ and $b$ are two coefficients. The total free energy of the system is

$$F(d,l) = E_m(d) + N_s\left[V(l) + E_s\left(x_v, l\right)\right]. \tag{17}$$

We sought to determine the equilibrium membrane distance and the corresponding SNARE folding state as a function of the tilt angle $\alpha$, given other model parameters, including $N_s, L, R, a, b, A_m, d_m,$ and $V(l)$.. To this end, we first calculated the total free energy $F(d,l)$ as a function of the membrane distance $d$ and the contour length $l$. Note that the extension of the unfolded VAMP2 polypeptide in Eqs. (15) and (17) is an implicit function of $d$ and $l$. We then minimized this energy function to determine the equilibrium membrane distance and contour length.

The extension of the unfolded VAMP2 $x_v$ depends on the position and configuration of the trans-SNARE complex between the PM and the vesicle membrane (Figures S16A–S16E). We defined the y-axis in the PM plane and the x-axis as passing through the center of the spherical vesicle. Given the membrane position and SNARE folding state, the extension $x_v$ depends on the position of the trans-SNARE complex, which can be described by the y-coordinate of point A, designated $y$ (Figure S16A). Accordingly, the positions are point A at $(0, y)$, point B at $(L\sin\alpha, y + L\cos\alpha)$, point D at $(h\sin\alpha, y + h\cos\alpha)$, and point O at $(d + R, 0)$. The extension of the unfolded VAMP2 $x_v$ can be calculated as

$$x_v(l, d, y) = |DO| - |EO| = \sqrt{(h\sin\alpha - R - d)^2 + (y + h\cos\alpha)^2} - R. \tag{18}$$

We then minimized the energy $x_v$ with respect to the SNARE position $y$, which depends on the membrane distance. If the membrane distance is greater than the height of the SNARE rod (Figure S16B), or

$$d > d_1 = L\sin\alpha, \tag{19}$$

the SNARE complex can move freely in the PM without touching the vesicle and has a minimum extension or total energy with

$$x_v(l, d, y_{\min 1}) = d - h\sin\alpha \tag{20}$$

when

$$y_{\min 1} = -h\cos\alpha. \tag{21}$$

When $d \le d_1$,, the SNARE rod will eventually touch the SUV at either the N-terminal point (Figure S16C) or an internal point (Figure S16D), as the trans-SNARE complex moves toward the SUV. The latter is a point of tangency on the vesicle, with a critical SNARE length $L_c = |AB| \le L$ (Figure S16E). The contact site is determined by the following equations

$$\tan\beta = \frac{R}{L_c - \dfrac{d}{\sin\alpha}}, \tag{22}$$

where

$$\alpha = \pi - 2\beta. \tag{23}$$

Therefore,

$$L_c = R\tan\left(\frac{\alpha}{2}\right) + \frac{d}{\sin\alpha}. \tag{24}$$

The tangent contact occurs when the membrane distance is less than a critical value $d_2$, with

$$d < d_2 \equiv \left[L - R\tan\left(\frac{\alpha}{2}\right)\right]\sin\alpha. \tag{25}$$

The tangent contact also requires $d_2 \ge 0$, or

$$L \geq R\tan\left(\frac{\alpha}{2}\right). \tag{26}$$

Therefore, the SNARE rod can touch the SUV when $d \leq d_1$ and can contact it tangentially at an internal site when $0 \leq d < \max(d_2, 0)$. For the calculations below, we define the minimum of $L$ and $L_c$ as $L_{eff}$, or $L_{eff} = \min(L, L_c)$. When the SNARE rod touches the vesicle, the SNARE position has a minimum value of $y = y_{\min 2}$. Because the contact site is located on the circle, we have

$$\begin{aligned} &\left(L_{eff}\sin\alpha - R - d\right)^2 + \left(y_{\min 2} + L_{eff}\cos\alpha\right)^2 = R^2 \\ &y_{\min 2} = \sqrt{R^2 - \left(L_{eff}\sin\alpha - R - d\right)^2} - L_{eff}\cos\alpha \end{aligned} \tag{27}$$

Substitution of Eq. (27) into Eq. (18) yields the extension $x_v(l, d, y_{\min 2})$ when the SNARE rod touches the SUV. Therefore,

$$x_v(l,d) = \begin{cases} d - h\sin\alpha, \quad d \geq d_1; \\ \min\left(x_v(l,d,y_{\min 1}), x_v(l,d,y_{\min 2})\right), \quad d < d_1, y_{\min 1} > y_{\min 2}; \\ x_v(l,d,y_{\min 2}), \quad d < d_1, y_{\min 1} \leq y_{\min 2}. \end{cases} \tag{28}$$

For $\alpha > 0$, SNARE zippering cannot be complete in our model because neither the SNARE rod nor the membranes bend. The minimum SNARE unfolding $l_{\min}$ can be derived by setting $d = 0$ and $x_v(l, d, y) = l$ in Eqs. (18) and (24) and noting Eq. (16) (Figure S16E). Specifically, $l_{\min}$ is the solution to the following quadratic function with respect to $l$:

$$\left(1 - a^2\right)l^2 + 2\left(Ra\sin\alpha - ya\cos\alpha + R - ab\right)l - y^2 - b^2 + 2Rb\sin\alpha - 2yb\cos\alpha = 0, \tag{29}$$

where

$$\begin{aligned} &y = \sqrt{R^2 - \left(L_{eff}\sin\alpha - R\right)^2} - L_{eff}\cos\alpha, \\ &L_{eff} = \min(L, L_c), \\ &L_c = R\tan\left(\frac{\alpha}{2}\right). \end{aligned} \tag{30}$$

Numerical calculations showed that the minimum required VAMP2 unfolding ($l_{\min}$) is 0.05, 7.1, and 15.1 residues for $\alpha = 5°$, 60°, and 90°, respectively, with $R = 20$ nm, $a = 0.15/0.365 = 0.41$, and $b = 0$ (Figure S16F).

Finally, the equilibrium distance between the vesicle and PM and the conformation of the trans-SNARE complex can be obtained by minimizing $F(d, l)$ with respect to $d$ and $l$, or

$$\begin{cases} \dfrac{\partial F}{\partial d} = 0 \\ \dfrac{\partial F}{\partial l} = 0. \end{cases} \tag{31}$$

The computational procedure is summarized below:

1. Define the model parameters $L, R, a, b, A_m, d_m, P, N$ and the SNARE folding energy landscape $V(l)$.
2. Given $d$ and $l$, calculate the critical membrane distance $d_1 = L\sin\alpha$, $y_{\min 1} = -h\cos\alpha$, $y_{\min 2}$, the critical SNARE length $L_c = R\tan\left(\frac{\alpha}{2}\right) + \frac{d}{\sin\alpha}$, and the effective SNARE length $L_{eff} = \min(L, L_c)$.
3. Calculate the VAMP2 extension $x_v(l,d)$ according to Eq. (28).
4. Calculate the VAMP2 stretching energy $E_s(x_v(l,d),l)$ according to Eq. (15), the stretch force $F$ according to Eq. (14), the membrane repulsive energy $E_m(d)$ according to Eq. (11), the SNARE folding energy $V(l)$ based on the provided energy function, and the total free energy function $F(d,l)$ based on Eq. (17)
5. Given $d$, minimize $F(d,l)$ with respect to $l$, yielding the minimum energy function $F_{\min l}(d)$ and the optimal contour length $l_e(d)$, which gives the equilibrium SNARE folding state as a function of membrane distance.
6. Given $l$, minimize $F(d,l)$ with respect to $d$, yielding the minimum energy function $F_{\min d}(l)$ and the optimal membrane distance $d_e(l)$, which gives the equilibrium membrane distance as a function of SNARE folding state.

**T6. Determination of Gβγ-dependent SNARE complex clamping kinetics**

Because Gβγ-clamped states were long lived, constant-force extension trajectories were often recorded for up to 1 h and sometimes exhibited substantial baseline drift. We used two complementary analysis approaches. Trajectories with limited baseline drift were analyzed by hidden Markov modeling (HMM), using a three-state model when NTD unzipping was absent and a four-state model when it was present. These analyses yielded clamping and de-clamping rates directly. Trajectories with substantial drift were analyzed by manual identification of long-lived Gβγ-clamped intervals. For these trajectories, a dwell-time threshold of 0.5 s defined a Gβγ-clamped state. Clamping rates were then calculated from the rate expression derived below.

Let the probabilities that the SNARE complex occupies the zippered state (F in the diagram below), the unzipped or half-zippered state (U), and the Gβγ-clamped state (B) be $p_F$, $p_U$, and $p_B$, respectively. Given the transition rates shown in the diagram, the state probabilities satisfy the following equations under equilibrium conditions:

$$F \underset{k_f}{\overset{k_u}{\rightleftharpoons}} U$$

$$U \underset{k_d}{\overset{k_b = k_{on}[P]}{\rightleftharpoons}} B$$

$$\begin{cases} p_F k_u = p_U k_f \\ p_U k_b = p_B k_d \\ p_F + p_U + p_B = 1. \end{cases} \tag{32}$$

Here $k_u$ is the unfolding rate of the zippered state, $k_f$ is the folding rate of the half-zippered state, $k_b$ is the rate of Gβγ clamping of the half-zippered state, and $k_d$ is the de-clamping rate. Suppose the total numbers of visits to the three states are $N_F$, $N_U$, and $N_B$ during a constant-force extension trajectory of total duration $T$. Then

$$p_i = \frac{N_i \tau_i}{T},\ i = F, U, B, \tag{33}$$

where $\tau_i$ is the average lifetime of state $i$. For the reaction scheme shown above, the average state lifetimes are related to the rate constants as

$$\begin{cases} \tau_F = \dfrac{1}{k_u}, \\ \tau_U = \dfrac{1}{k_f + k_b}, \\ \tau_B = \dfrac{1}{k_d}. \end{cases} \tag{34}$$

Substitution of Eqs. (33) and (34) into Eq. (32) yields

$$\begin{cases} N_F = \dfrac{N_U k_f}{k_f + k_b} \\ \dfrac{N_U k_b}{k_f + k_b} = N_B \\ \dfrac{N_F}{k_u} + \dfrac{N_U}{k_f + k_b} + \dfrac{N_B}{k_d} = T \end{cases} \tag{35}$$

We solved for $k_b$ from Eq. (35), yielding

$$k_b = \frac{N_B}{p\left(T - T_B\right)}, \tag{36}$$

where $T_B = N_B \tau_B$ is the total time spent in the Gβγ-clamped state, and $p$ is the fraction of time outside the Gβγ-clamped state that the SNARE complex spends in the half-zippered state, with

$$p = \frac{p_U}{p_F + p_U}. \tag{37}$$

Note that $p\left(T - T_B\right)$ is the total time that the unclamped SNARE complex spends in the half-zippered state.

Numerical calculations, including hidden Markov modeling, energy-landscape fitting, and simulations, were performed in MATLAB. Custom code will be made available before publication.

### Supplementary Figures

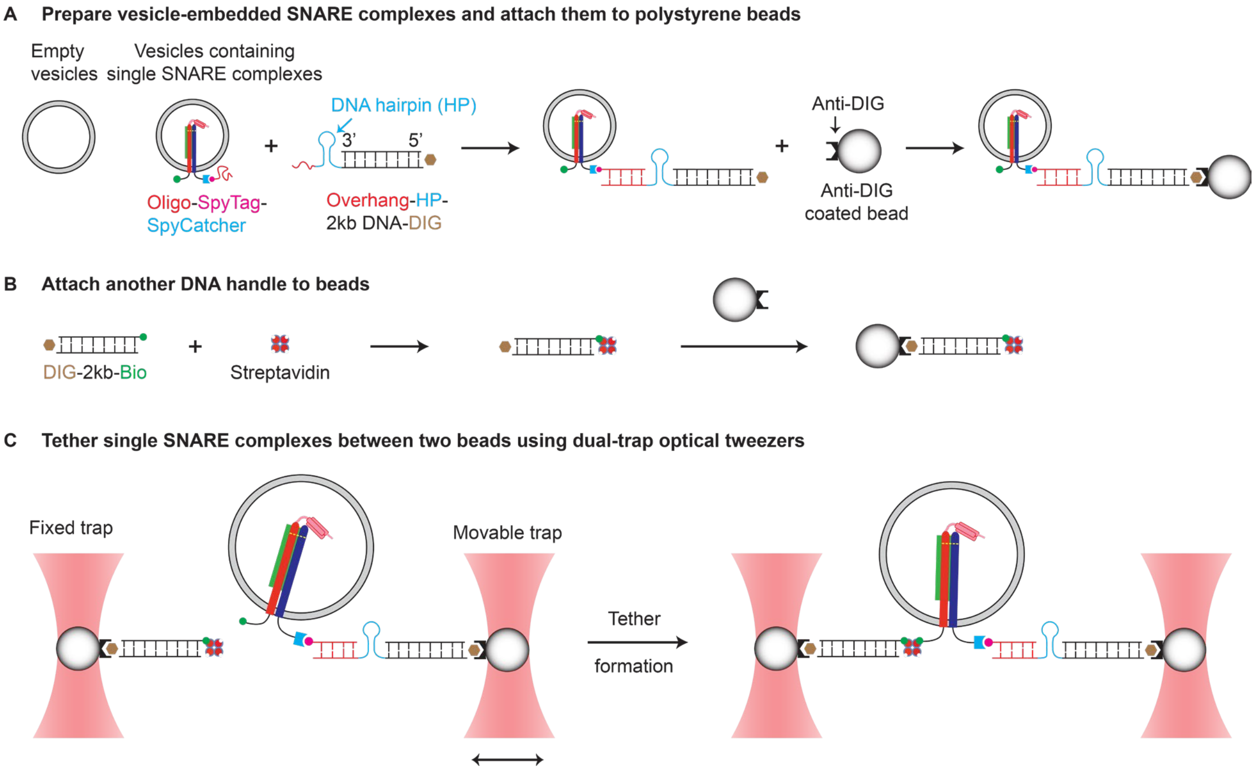


**Figure S1. Assembly of single cis-SNARE complexes for single-molecule optical tweezers measurements.**

(A) Proteoliposomes containing SNARE complexes were incubated with DNA handles bearing a single-stranded overhang that hybridized to a complementary oligonucleotide conjugated to VAMP2 through a SpyTag/SpyCatcher covalent linkage[6]. The DNA handle contained two digoxigenin (DIG) moieties at its 5′ end for subsequent bead attachment.

(B) A second DNA handle was coupled to tetrameric streptavidin through biotin and attached to an anti-digoxigenin-coated polystyrene bead.

(C) Beads from separate microfluidic channels were optically trapped and brought together to form a single protein–DNA tether in a dumbbell geometry for force-spectroscopy measurements.

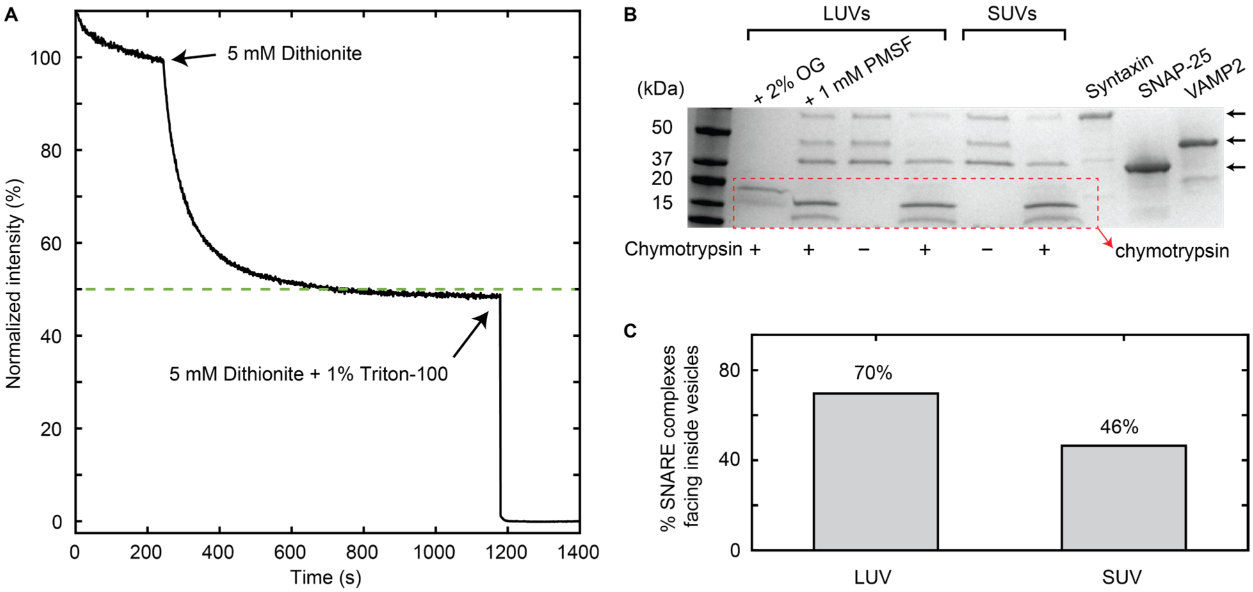


**Figure S2. Unilamellarity of SNARE-containing liposomes and orientation of reconstituted SNARE complexes.**

(A) Dithionite-quenching assay showing fluorescence loss after addition of dithionite and subsequent Triton X-100. The approximately 50% fluorescence decrease after dithionite addition is consistent with SUV unilamellarity.

(B) Chymotrypsin-protection assay showing selective digestion of SNARE proteins exposed on the vesicle exterior, whereas lumen-facing populations remained protected. Positive (SUV + protease + 2% OG) and negative (SUV + protease + 1 mM PMSF) controls validated complete and minimal digestion, respectively.

(C) Percentage of SNARE complexes facing the vesicle interior, determined by the protease-protection assay.

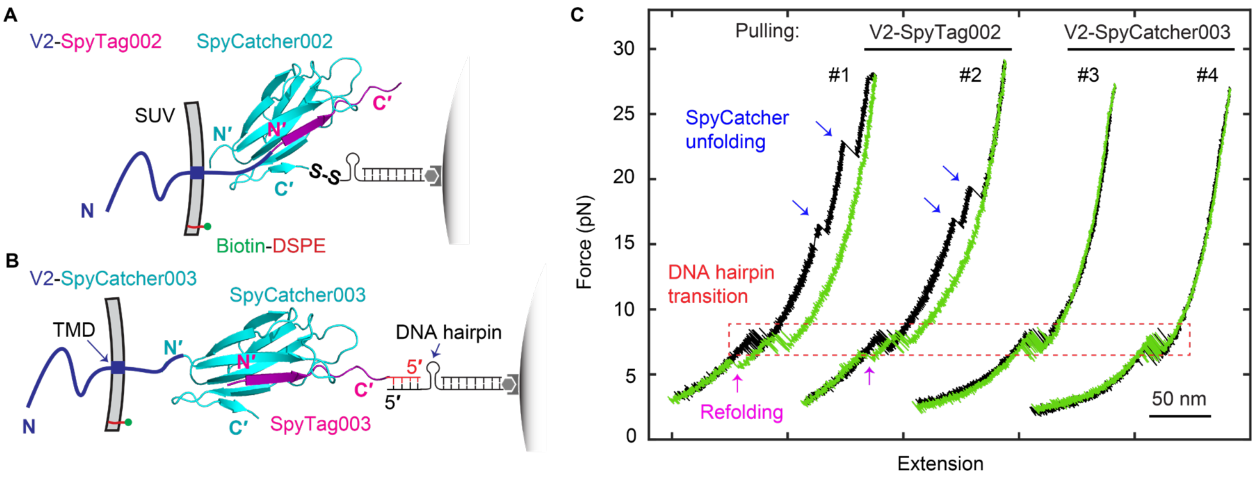


**Figure S3. Two SpyTag–SpyCatcher linkages used to pull cis-SNARE complexes.**
(A) Schematic of the VAMP2 (V2)-to-DNA linkage used in early cis-SNARE pulling experiments. SpyTag002 was fused to the C terminus of membrane-embedded VAMP2 and conjugated to SpyCatcher002 (PDB ID: 4MLI).[6] The DNA handle was crosslinked to the C terminus of SpyCatcher002 through a disulfide bridge. This pulling site produced SpyCatcher unfolding and refolding events in some force-extension curves (FECs)[7], including those in Figures 1C and S11. N and N′ mark the N termini of VAMP2 and SpyCatcher or SpyTag, respectively.
(B) Improved VAMP2-to-DNA linkage used in most SNARE pulling experiments. SpyCatcher003 was fused to membrane-embedded VAMP2 and conjugated to a chemically synthesized SpyTag003-oligonucleotide. The oligonucleotide hybridized to the overhang sequence of the DNA handle. The covalent VAMP2-DNA linkage through SpyTag003/SpyCatcher003 was mechanically stable over the force range used to pull SNARE complexes.
(C) Representative FECs obtained by pulling single SUV-anchored VAMP2 molecules using the two configurations shown in (A) and (B). Single VAMP2 molecules were pulled from the VAMP2 C terminus and biotin-DSPE in SUV membranes. FECs #1 and #2 display multiple discrete unfolding rips (blue arrows), corresponding to mechanical unfolding of SpyCatcher002 domains, followed by refolding events during relaxation (magenta arrows). By contrast, FECs #3 and #4 lack protein unfolding rips and exhibit smooth, continuous force-extension curves. A DNA hairpin connected in series to VAMP2 yielded the reversible DNA-hairpin unfolding/refolding transitions marked by the dashed red rectangle.

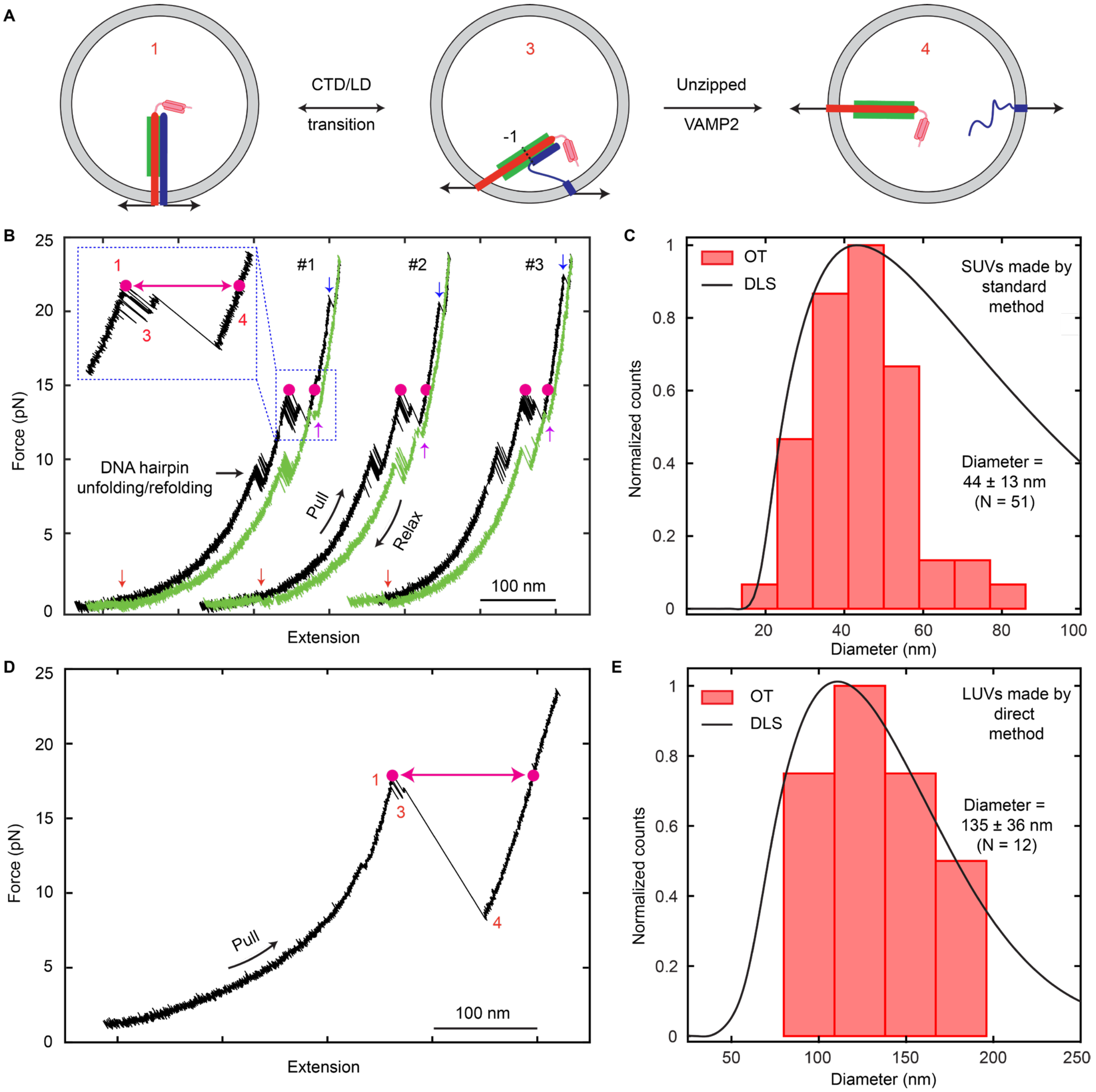


**Figure S4. Vesicle sizes measured by optical tweezers and dynamic light scattering.**
(A) SNARE states obtained by pulling single uncrosslinked cis-SNARE complexes to detect both SNARE folding/unfolding transitions and vesicle size. Upon pulling to high force, the cis-SNARE complex undergoes sequential transitions from (1) an initially assembled state to (2) a half-zippered state and then (3) an unzipped state. Transition to the unzipped state is accompanied by an extension jump limited by the vesicle diameter.
(B) Representative force-extension curves (FECs) showing multiple unfolding and refolding cycles of a single uncrosslinked cis-SNARE complex. Individual pulling cycles (#1 to #3) are offset along the extension axis for clarity. Magenta point pairs indicate extension changes approximately equal to the vesicle diameter. The inset shows an enlarged view of the boxed CTD/LD and NTD transition region in FEC #1. Blue and magenta arrows denote force-induced

unfolding and refolding, respectively, of SpyCatcher002 conjugated to VAMP2 (Figure S3A). Red arrows denote reassembly of the SNARE complex at low force.

(C) Distributions of SUV diameters derived from optical-tweezers measurements (red bars) and dynamic light scattering (DLS; black curve). The average diameters measured by the two approaches are consistent: 44 ± 13 nm (mean ± standard deviation) by optical tweezers and 43 ± 3 nm by DLS.

(D) Representative FEC obtained by pulling cis-SNARE complexes reconstituted on LUVs.

(E) Distributions of LUV diameters derived from optical-tweezers measurements (red bars) and DLS (black curve). The average diameters measured by the two approaches are consistent within experimental variability: 135 ± 36 nm by optical tweezers and 110 ± 8 nm by DLS. The slightly larger LUV size measured by optical tweezers likely reflects LUV membrane elongation in the pulling direction.

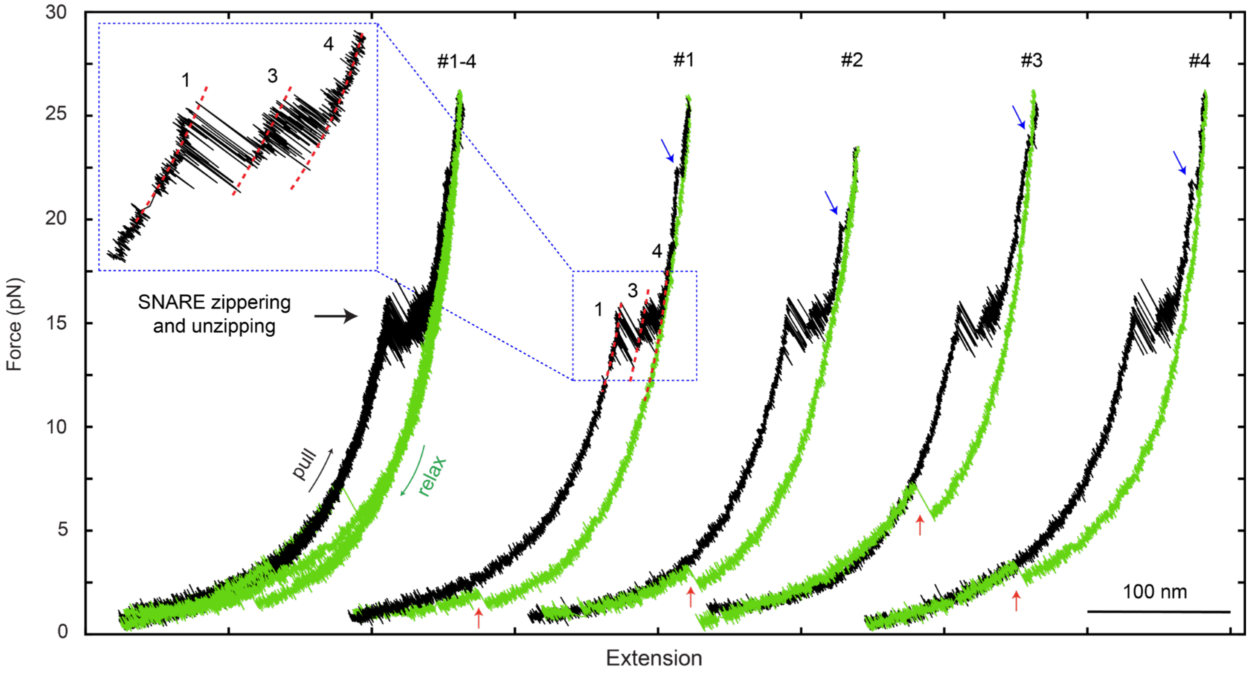


**Figure S5. Force-extension curves (FECs) of a single cis-SNARE complex without a DNA hairpin connected in series.** Overlapping FECs from repeated pulling–relaxation cycles (#1 to #4) of a single cis-SNARE complex on a pure POPC SUV show characteristic three-state unfolding and refolding transitions and full reassembly of the SNARE complex at low force. Removal of the DNA hairpin did not reveal additional mechanical intermediates. The blue and red arrows show t-SNARE unfolding and SNARE complex reassembly, respectively. The inset shows an expanded view of the zippering region from pull #1, revealing reversible three-state transitions.

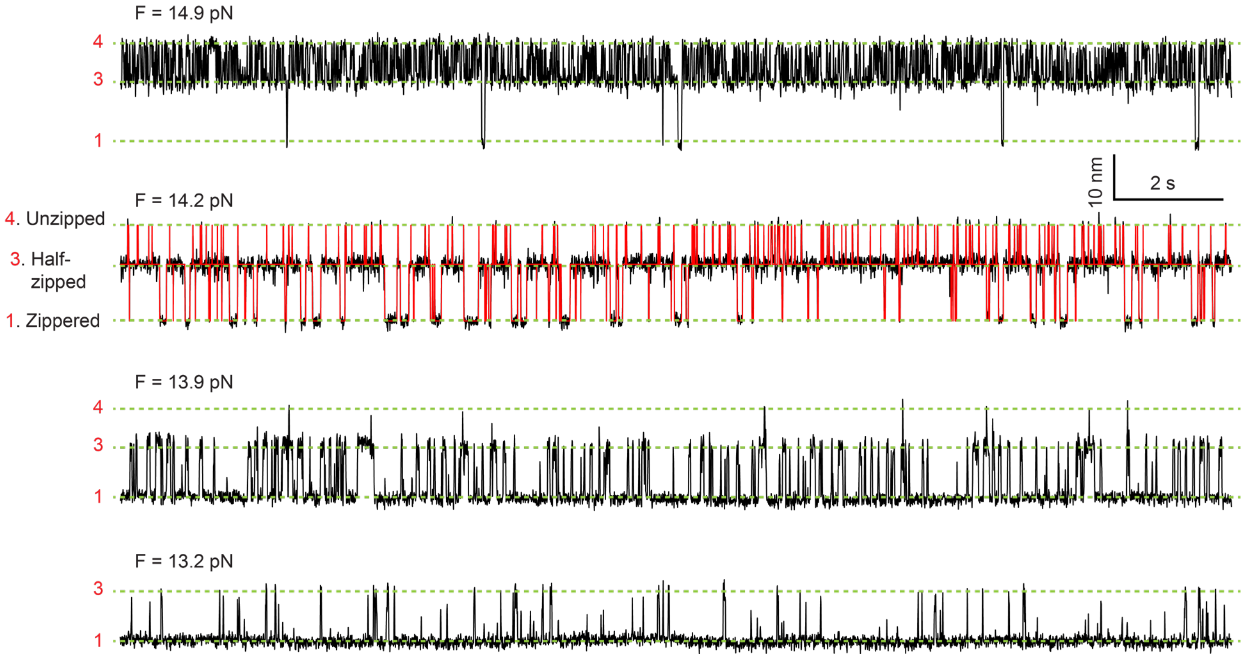


**Figure S6. Force-dependent extension trajectories of a single cis-SNARE complex on a POPC SUV.** Representative trajectories at the indicated constant mean forces (F) show three-state SNARE folding/unfolding transitions. Distinct extension states corresponding to fully zippered, half-zippered, and unzipped configurations are indicated by green dashed lines and labeled as states 1, 3, and 4, respectively (Figure 1E).

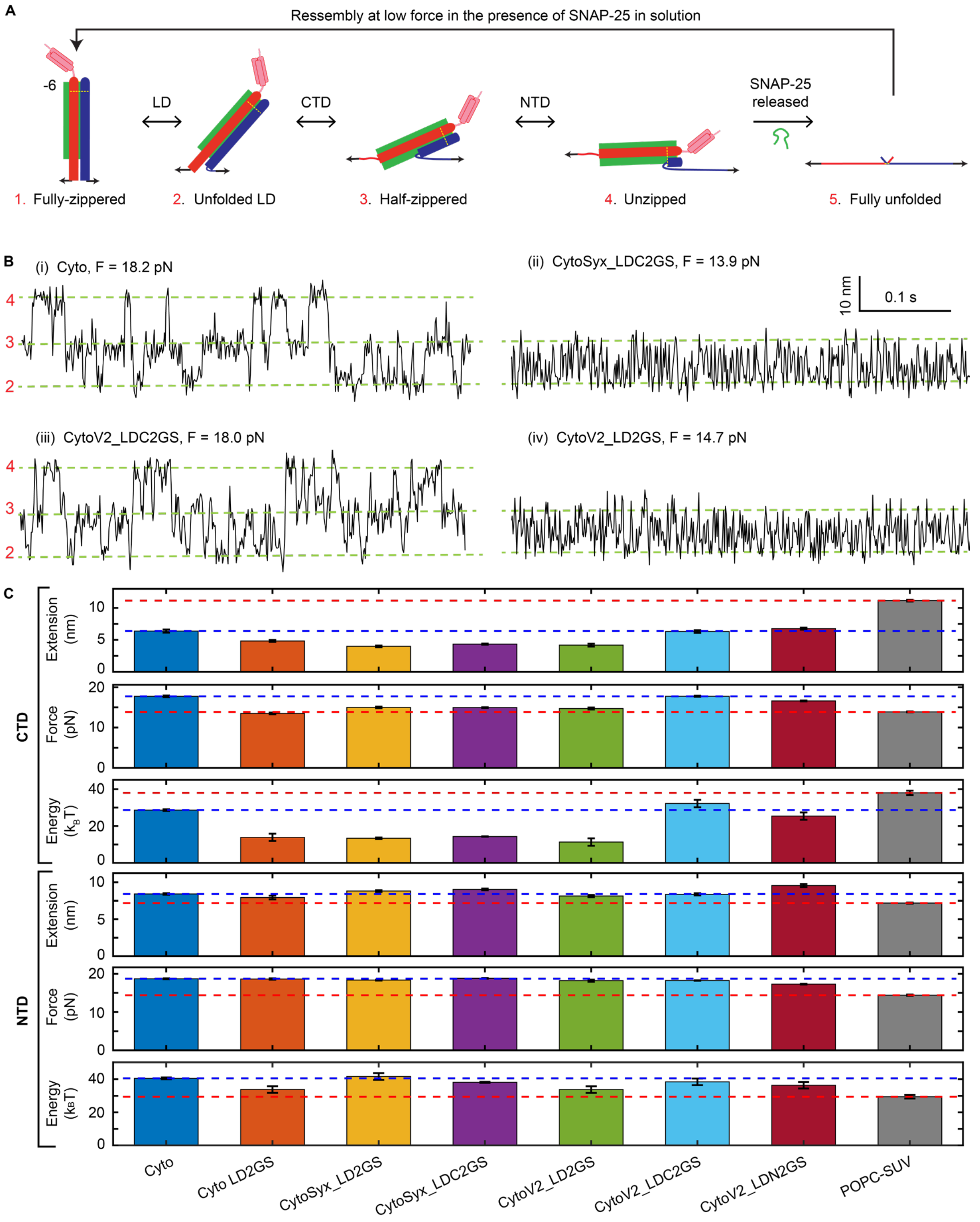


**Figure S7. Folding intermediates and energetics of cytosolic SNARE complexes and linker-domain variants.**

(A) Revised model for cytosolic SNARE folding and unfolding. In LD-unzipped state 2, the syntaxin LD remains helical, whereas VAMP2 is unfolded to residue 89 ± 1 (mean ± SEM). This asymmetric state differs from our previous model, in which both the syntaxin and VAMP2 LDs were proposed to unfold[8-10]. The data in (B) support the revised structural assignments.

(B) Representative constant-force extension trajectories of the WT cytosolic SNARE complex and variants containing LD replacements in syntaxin or VAMP2 (Table S1). Data were filtered to 500 Hz. In Syx_LDC2GS, the C-terminal five residues of the syntaxin LD (261 to 265) were replaced by a GS linker (Figure 2D). In V2_LD2GS, the entire VAMP2 LD sequence was replaced by a GS linker. Both substitutions destabilized CTD zippering relative to WT, as indicated by lower unzipping forces (~15 pN), suggesting that the corresponding LD regions are at least partially folded in the half-zippered state. By contrast, replacing the C-terminal portion of the VAMP2 LD in V2_LDC2GS did not affect CTD zippering, indicating that this segment is unfolded in the half-zippered state. Because LD unfolding occurs at an equilibrium force of ~12 pN[8,9], these results further suggest that LD unfolding involves asymmetric unzipping of VAMP2 from a helical syntaxin LD.

(C) Mean extension changes, equilibrium forces, and free energies for the CTD and NTD folding–unfolding transitions of the indicated cytosolic SNARE complexes, LD mutants, and WT cis-SNARE complex. Blue and red dashed horizontal lines indicate the corresponding values for the WT cytosolic SNARE complex and WT cis-SNARE complex on POPC SUVs, respectively. Bars and error bars indicate mean ± SEM.

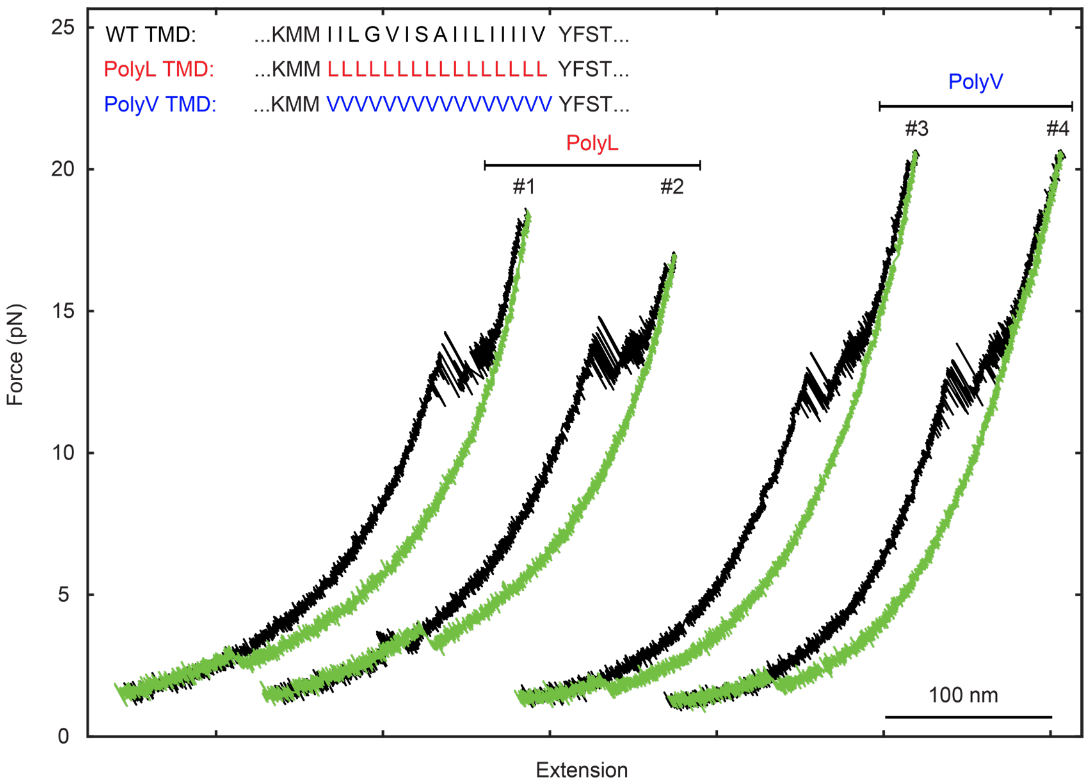


**Figure S8. SNARE TMDs exhibit little intrinsic affinity.** The VAMP2 TMD sequence was replaced with polyleucine (PolyL) or polyvaline (PolyV) sequences. These substitutions did not alter the force-extension curves compared with the WT construct, indicating little intrinsic affinity between the SNARE TMDs. VAMP2 TMD sequences for WT, PolyL, and PolyV constructs are shown in the inset.

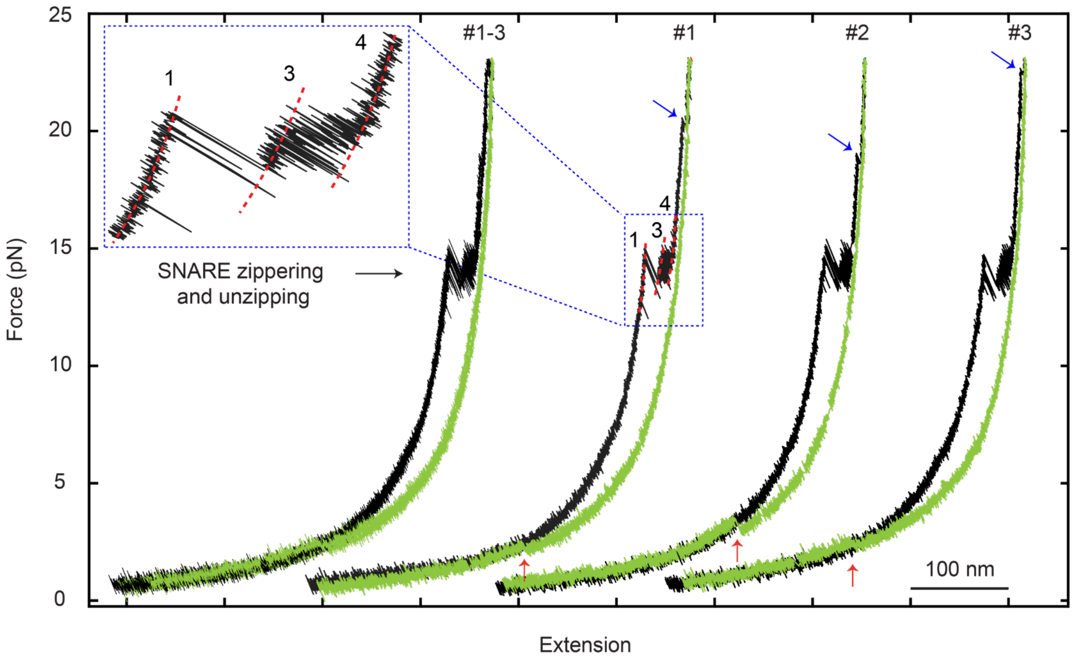


**Figure S9. Restoring native cysteine residues in TMDs does not alter cis-SNARE complex assembly.** Representative FECs (#1 to #3) of cis-SNARE complexes containing native cysteine residues in syntaxin and VAMP2 are shown. The observed unfolding and refolding transitions, as well as the overall force-extension curves, are indistinguishable from those measured for constructs in which the native TMD cysteines were replaced by serine (see also Figures 2A and S10). The inset shows an expanded view of the boxed zippering region in pulling curve #1.

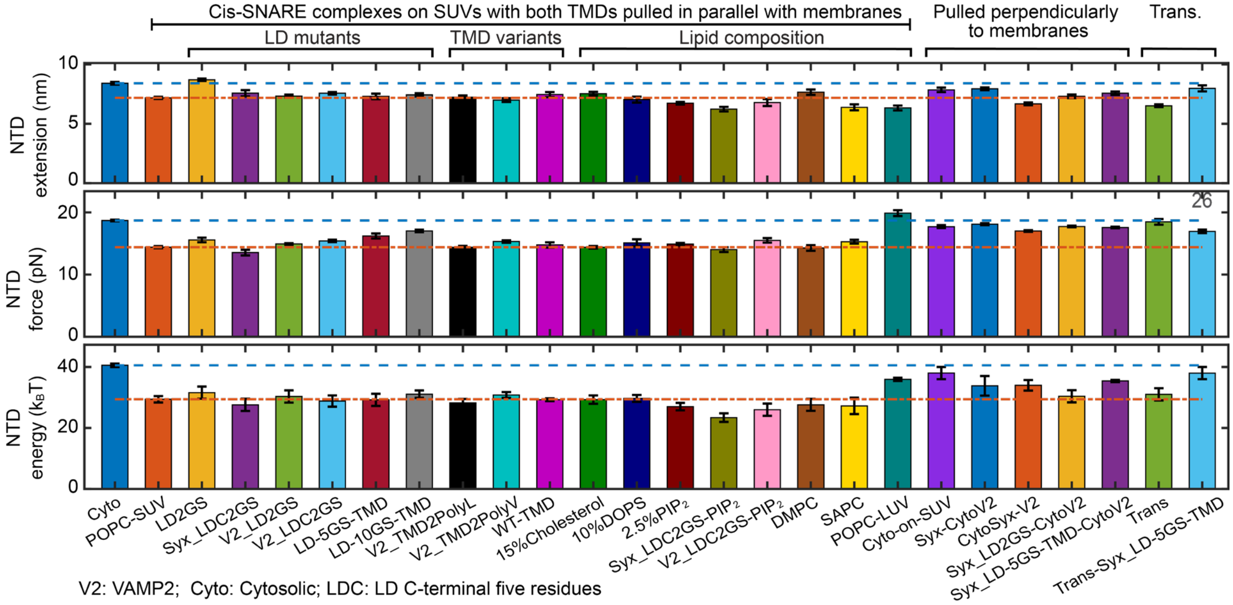


**Figure S10. Effects of SNARE domain variants and membrane environment on NTD folding mechanics.** Mean extension changes, equilibrium forces, and free energies for the NTD folding–

unfolding transition of the indicated SNARE complexes, LD mutants, TMD variants, lipid compositions, bilayer thicknesses, and membrane-anchoring configurations. Blue dotted and red dashed horizontal lines indicate the corresponding values for the WT cytosolic SNARE complex and WT cis-SNARE complex in 100 mol% POPC SUVs, respectively. Bars and error bars indicate mean ± SEM.

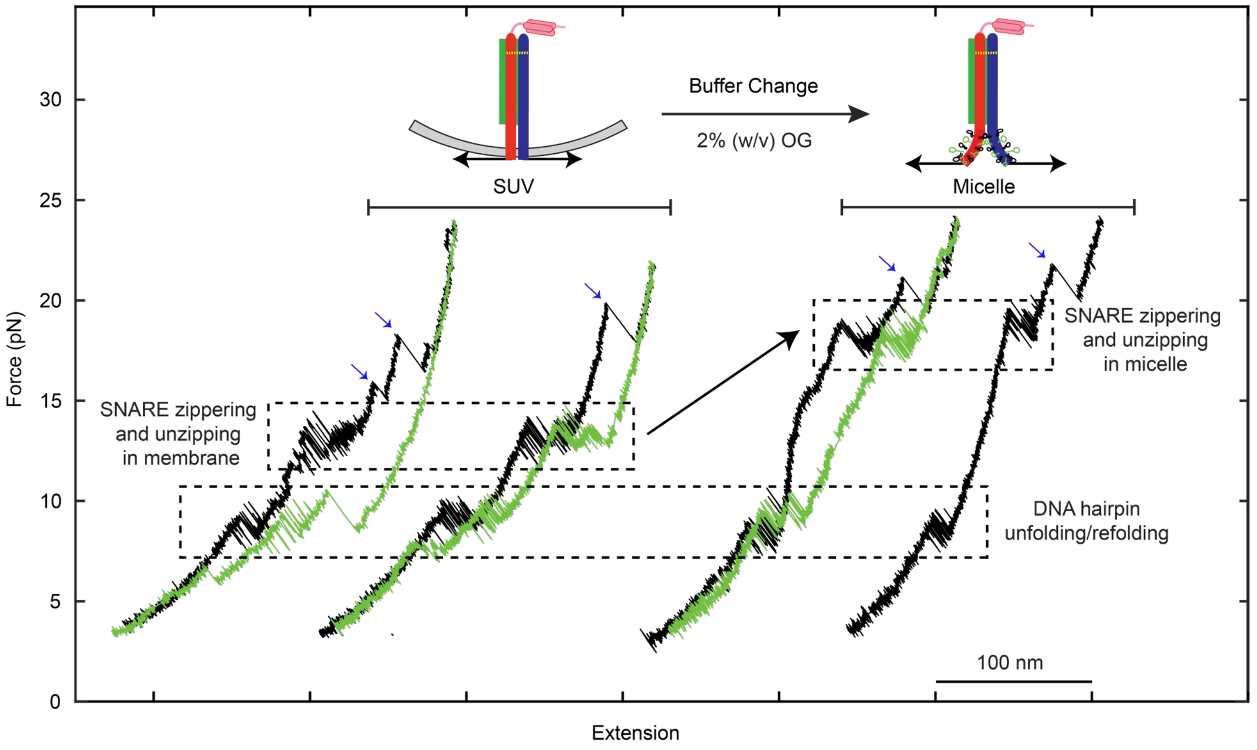


**Figure S11. SUV membranes reduce the SNARE zippering force.** Representative force-extension curves obtained by pulling a single SNARE complex first on an SUV and then in 2% (w/v) OG, showing reduced zippering force on bilayers. A single bilayer-embedded cis-SNARE complex was first pulled and relaxed for two rounds in a microfluidic channel to reveal its typical unfolding and refolding transitions. OG was then injected into the channel to dissolve the SUV and remove lipids from the SNARE proteins. The same SNARE complex was then pulled in OG micelles, leading to an upward shift in the SNARE transition force. Dashed boxes mark force ranges for reversible DNA-hairpin unfolding/refolding and SNARE zippering/unzipping, as indicated. Blue arrows mark SpyCatcher002 unfolding events (Figure S3A).

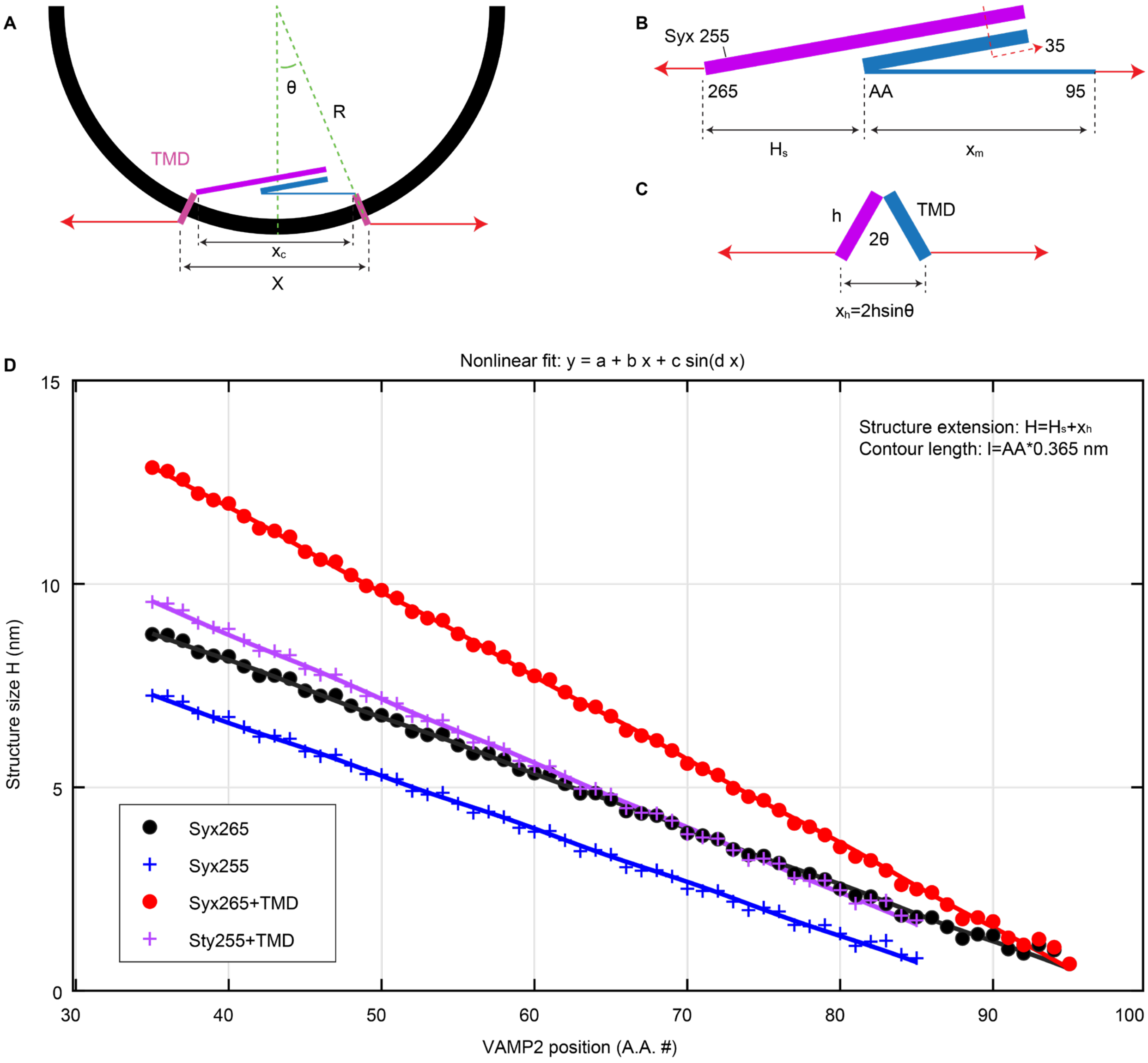


**Figure S12. Extension of the structured portion of the cis-SNARE complex along the pulling direction as a function of the last unfolded VAMP2 residue.**

(A) Diagram of a partially unfolded cis-SNARE complex inside a vesicle. The total SNARE extension along the pulling direction consists of contributions from structured and unfolded portions.

(B) The structured contribution includes the cytosolic SNARE bundle and TMD tilting see (C). The extension of the cytosolic SNARE bundle was defined as the distance from the last VAMP2 residue unfolded from the t-SNARE template to syntaxin residue Syx265 at the LD–TMD boundary, or Syx255 at the LD-CTD boundary, depending on the LD folding state. Distances were calculated from the crystal structure of the fully assembled SNARE complex (PDB ID: 3HD7), varying the last unfolded VAMP2 residue from 95 or 85 to residue 35 at the crosslinking site.

(C) Diagram of the extension change produced by TMD tilting.

(D) Extension of the structured portion and its nonlinear fit as a function of the last VAMP2 residue unfolded from the t-SNARE template. Extensions were calculated with (Syx265) or without

(Syx255) LD contributions and with (+TMD) or without TMD contributions. These values were used to fit measured extension changes and derive structures of folding intermediates (see Data analysis in Method details for more details).

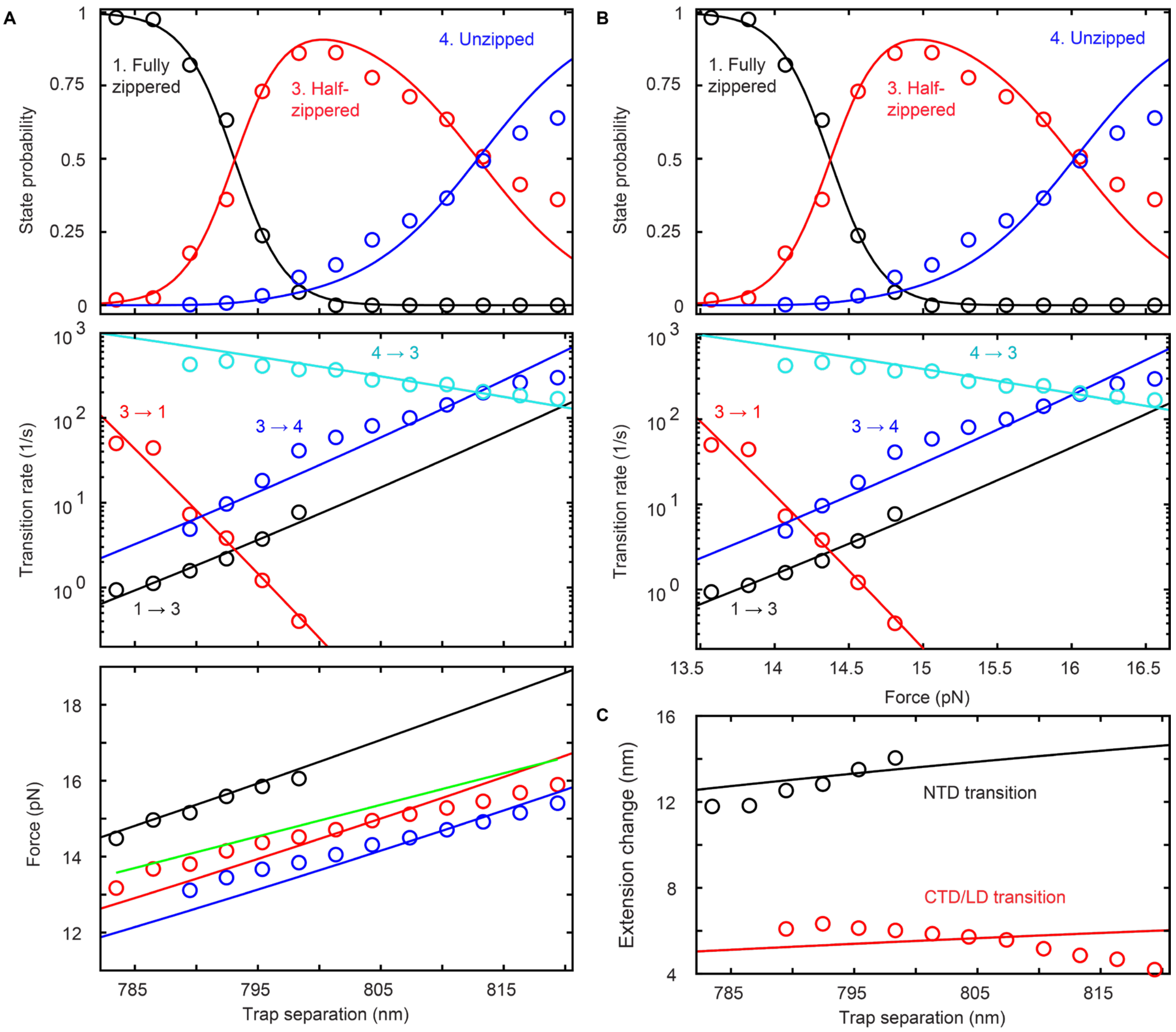


**Figure S13. Comparison of experimental measurements and best model fits used to determine the folding energy landscape of the cis-SNARE complex in the presence of $PIP_2$.**

(A) State probabilities (top), transition rates (middle), and average state forces (bottom) of the cis-SNARE complex as a function of trap separation. Symbols show measured values; solid lines show model fits, and states are color-coded as indicated. In the bottom panel, mean force as a function of trap separation was calculated as the average of the three best-fit state forces from linear fits to each state[11]. Experimental measurements were derived from hidden Markov modeling (HMM) of trap-separation-dependent extension trajectories.

(B) State probabilities (top) and transition rates (bottom) plotted as functions of mean force.

(C) Extension changes of the CTD/LD and NTD transitions as functions of trap separation. The best-fit model parameters yield the folding energy landscape of the cis-SNARE complex in the presence of 2.5 mol% $PIP_2$, as shown in Figure 1G.

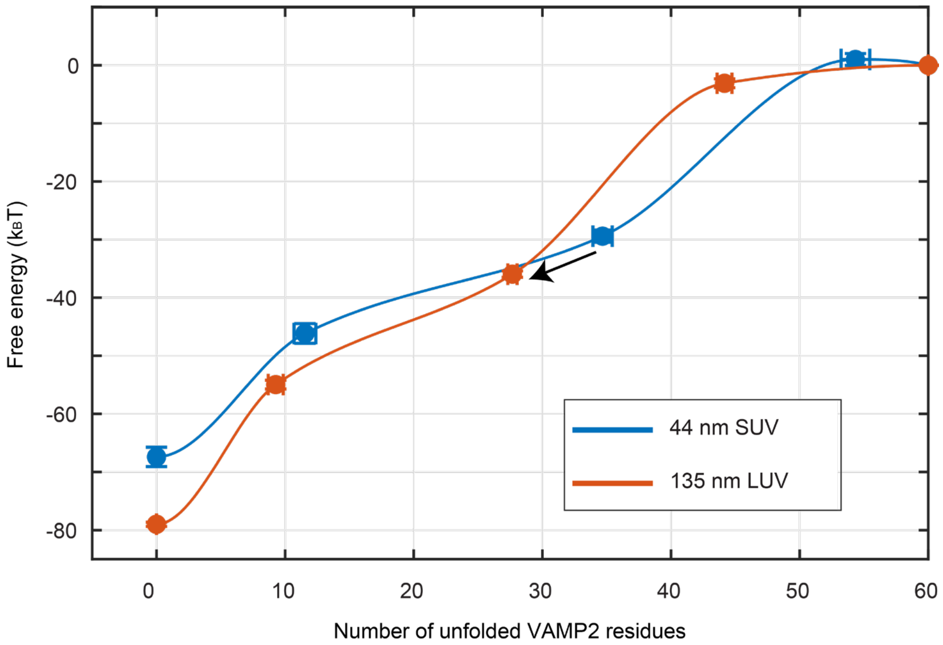


**Figure S14. Folding energy landscapes of the cis-SNARE complex on 44-nm SUVs and 135-nm LUVs.** The black arrow indicates the boundary shift of the half-zippered state. Symbols show experimental values; error bars indicate SEM; and lines are piecewise cubic fits to the experimental values.

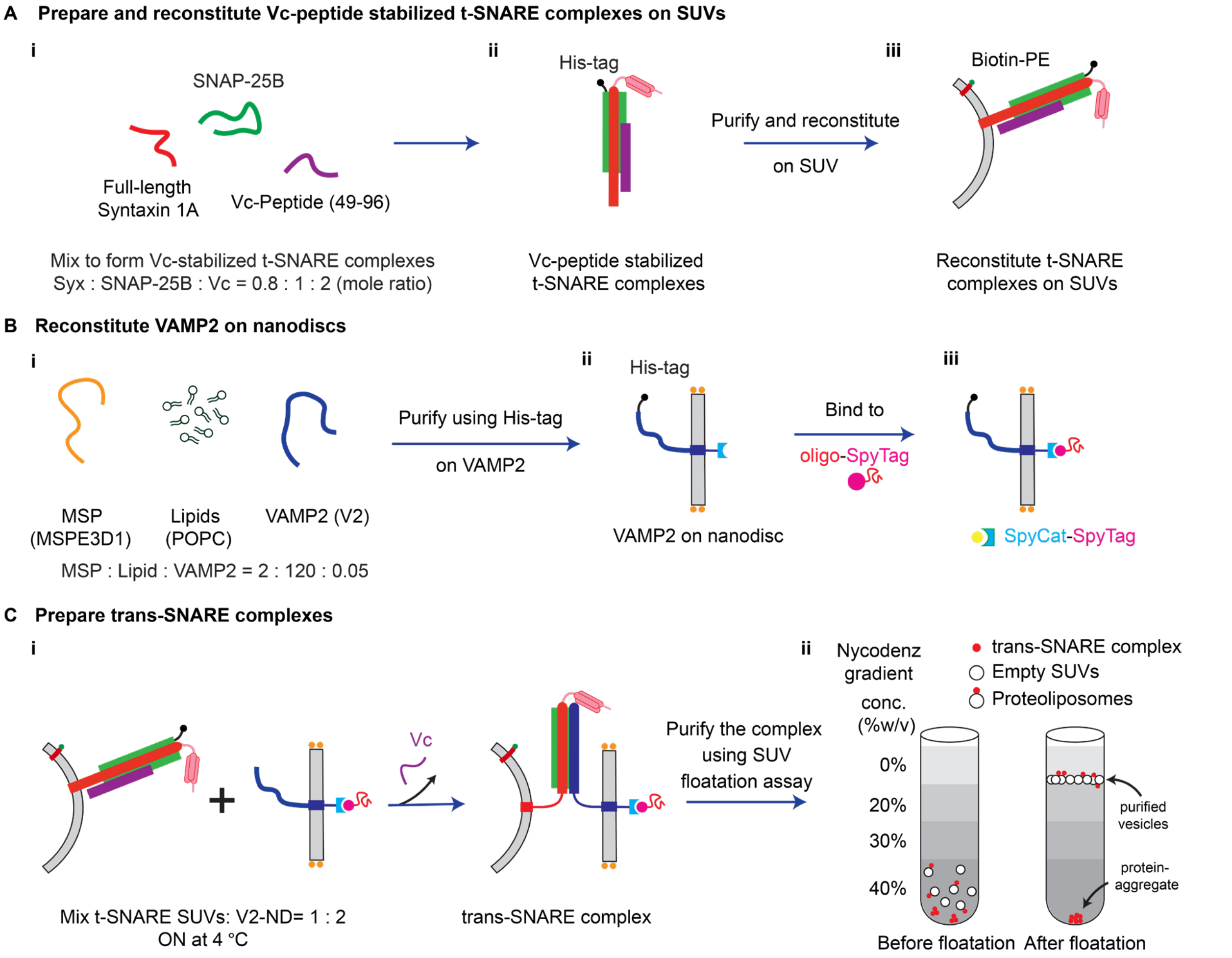


**Figure S15. Protocol for preparing trans-SNARE complexes for single-molecule pulling experiments.**

(A) Reconstitution of Vc-stabilized t-SNARE complexes into SUVs. (i) Full-length syntaxin, SNAP-25, and Vc peptide were mixed and incubated overnight to assemble Vc-stabilized t-SNARE complexes. Biotinylated lipids were included during SUV reconstitution to enable attachment to a DNA handle. (ii) Complexes were purified through the His tag on SNAP-25. (iii) Purified complexes were reconstituted into SUVs by OG-mediated co-micellization and dialysis.
(B) VAMP2 reconstitution in nanodiscs. (i) MSP, POPC, and VAMP2 were combined at defined molar ratios to assemble VAMP2-containing nanodiscs. (ii) Empty nanodiscs were removed by Ni-NTA affinity purification through the His tag on VAMP2. (iii) Purified VAMP2 nanodiscs were incubated with a two- to fivefold molar excess of SpyTag-oligo to enable downstream hybridization to DNA handles.
(C) Trans-SNARE assembly and purification. (i) t-SNARE-containing SUVs and VAMP2 nanodiscs were mixed and incubated overnight to form trans-SNARE complexes. The Vc peptide was displaced during trans-SNARE complex assembly[12]. (ii) Complexes were separated from unincorporated proteins and lipid aggregates by Nycodenz density-gradient flotation; SUVs migrated to the interface between the 0 and 10% Nycodenz layers.

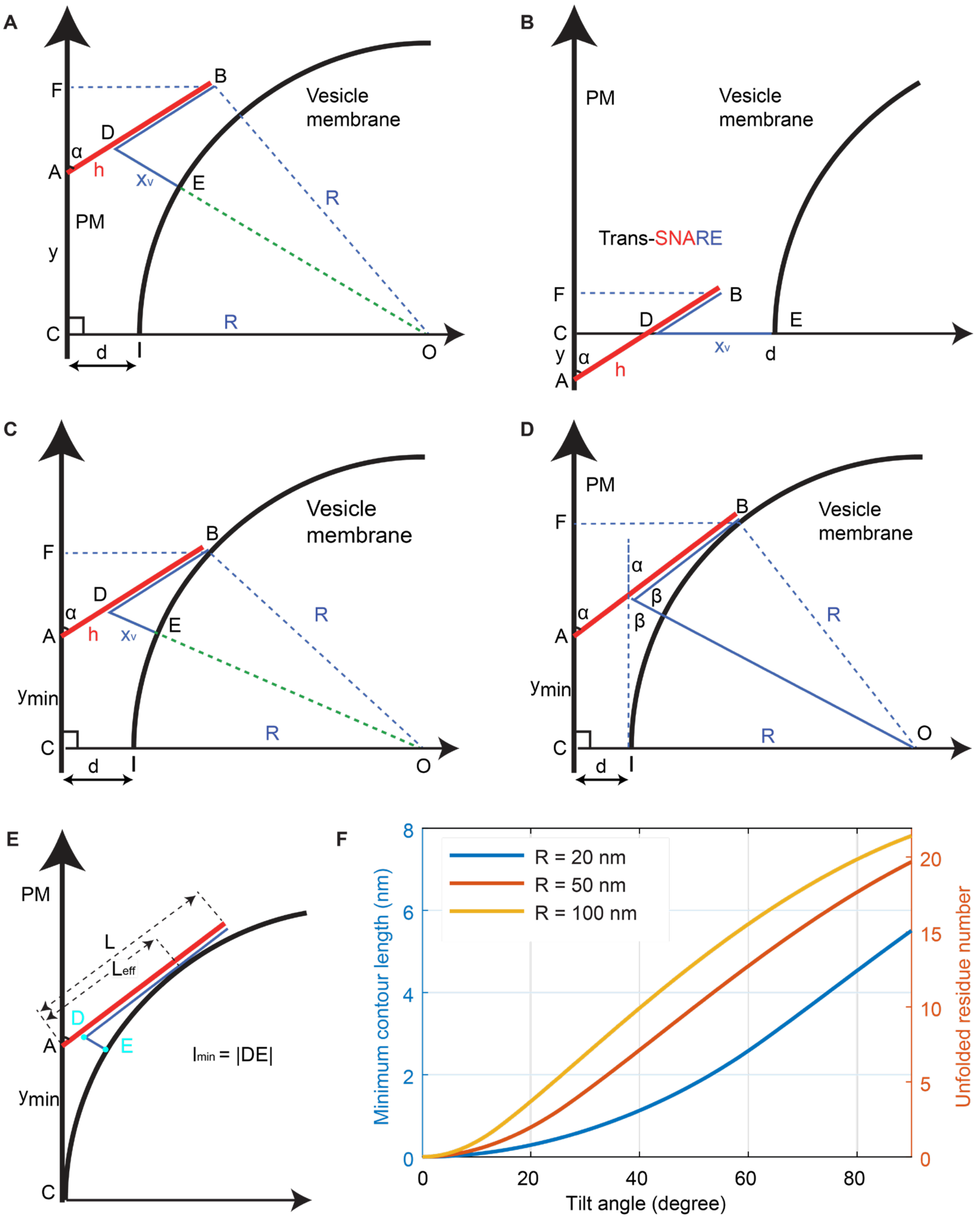


**Figure S16. Membrane-separation- and curvature-dependent conformations of trans-SNARE complexes with a fixed syntaxin tilt angle.**

(A) The partially zippered trans-SNARE complex is located at the vesicle edge without touching the vesicle membrane. At the indicated membrane separation, this complex remains at higher energy than the equilibrium state because shifting it downward lowers its energy (compare with the state in (C)).

(B) When membrane separation is sufficiently large, the partially zippered SNARE complex has minimum energy at the vesicle center.

(C) Representative equilibrium conformation for the trans-SNARE complex with minimum total free energy under the geometric constraints, in which the N terminus of the SNARE bundle touches the vesicle membrane.
(D) Another representative equilibrium conformation for a relatively small vesicle, in which the SNARE bundle begins to touch the vesicle membrane at an internal site.
(E) The SNARE bundle touches the vesicle membrane tangentially at an internal site.
(F) Minimum VAMP2 unfolding in the trans-SNARE complex as a function of syntaxin tilt angle for vesicles with different radii (R).

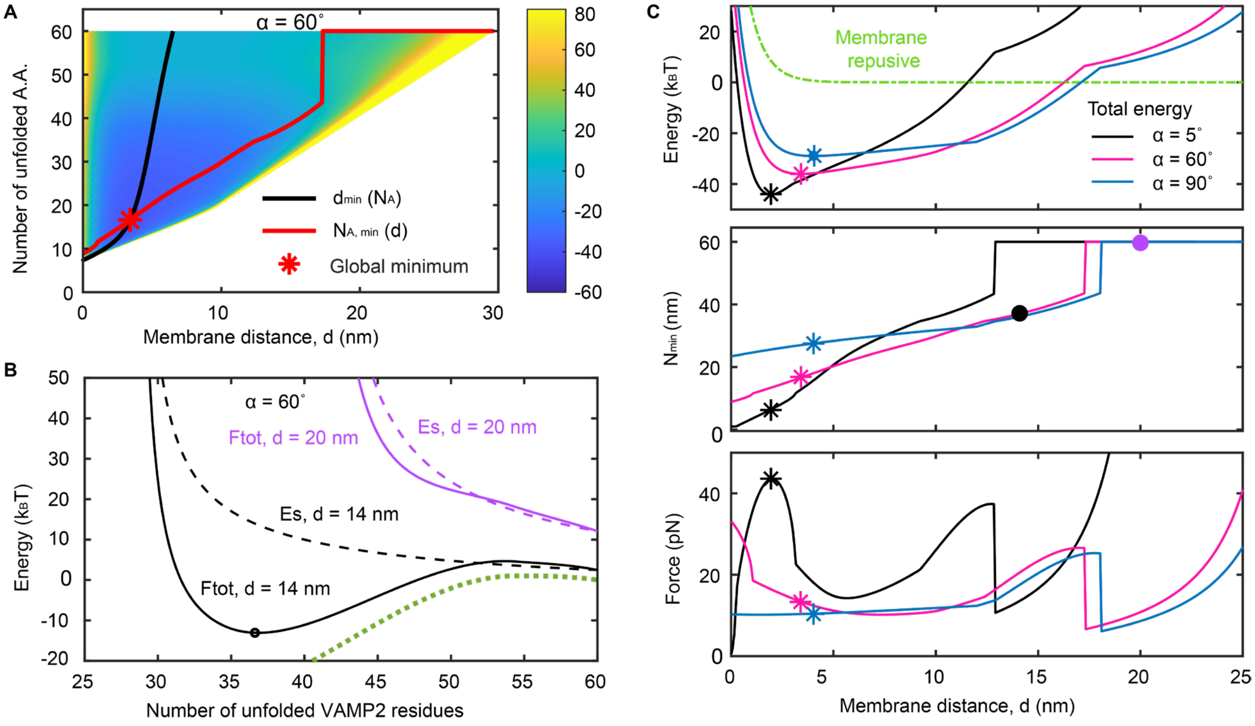


**Figure S17. Simulations showing that a large syntaxin tilt angle is required for efficient coupling of SNARE zippering to membrane apposition.**

(A) Contour map of the calculated total free energy of a system containing one trans-SNARE complex and two apposed membranes at a syntaxin tilt angle α = 60°, plotted as a function of membrane distance and the number of unfolded VAMP2 residues. The black curve shows the energy-minimizing membrane distance, and the red curve shows the energy-minimizing number of unfolded VAMP2 residues. The red star marks the global energy minimum.
(B) Energy as a function of the number of unfolded VAMP2 residues at membrane separations d = 14 nm and d = 20 nm. Solid curves, total free energy; dashed curves, entropic energy of the stretched unfolded VAMP2 polypeptide. Dotted curve, intrinsic energy landscape of the SNARE complex. The calculations show that at d = 20 nm, VAMP2 is completely unzipped, with the minimum energy located at the maximum modeled VAMP2 unfolding of 60 residues (constrained by the N-terminal VAMP2 crosslink). By contrast, at d = 14 nm, the energy has a minimum at approximately 37 unfolded residues, suggesting a half-zippered trans-SNARE complex. Thus, the SNARE complex begins to assemble only after membrane separation falls below a tilt-angle-dependent threshold. See also the middle panel in (C).
(C) Total free energy (top), equilibrium number of unfolded VAMP2 residues (middle), and SNARE stretching force (bottom) as functions of membrane separation at three syntaxin tilt angles.

Curves are color-coded by tilt angle. Stars mark global equilibrium values, and dots indicate equilibrium numbers of unfolded VAMP2 residues at the two membrane separations whose total energy is plotted in (B).

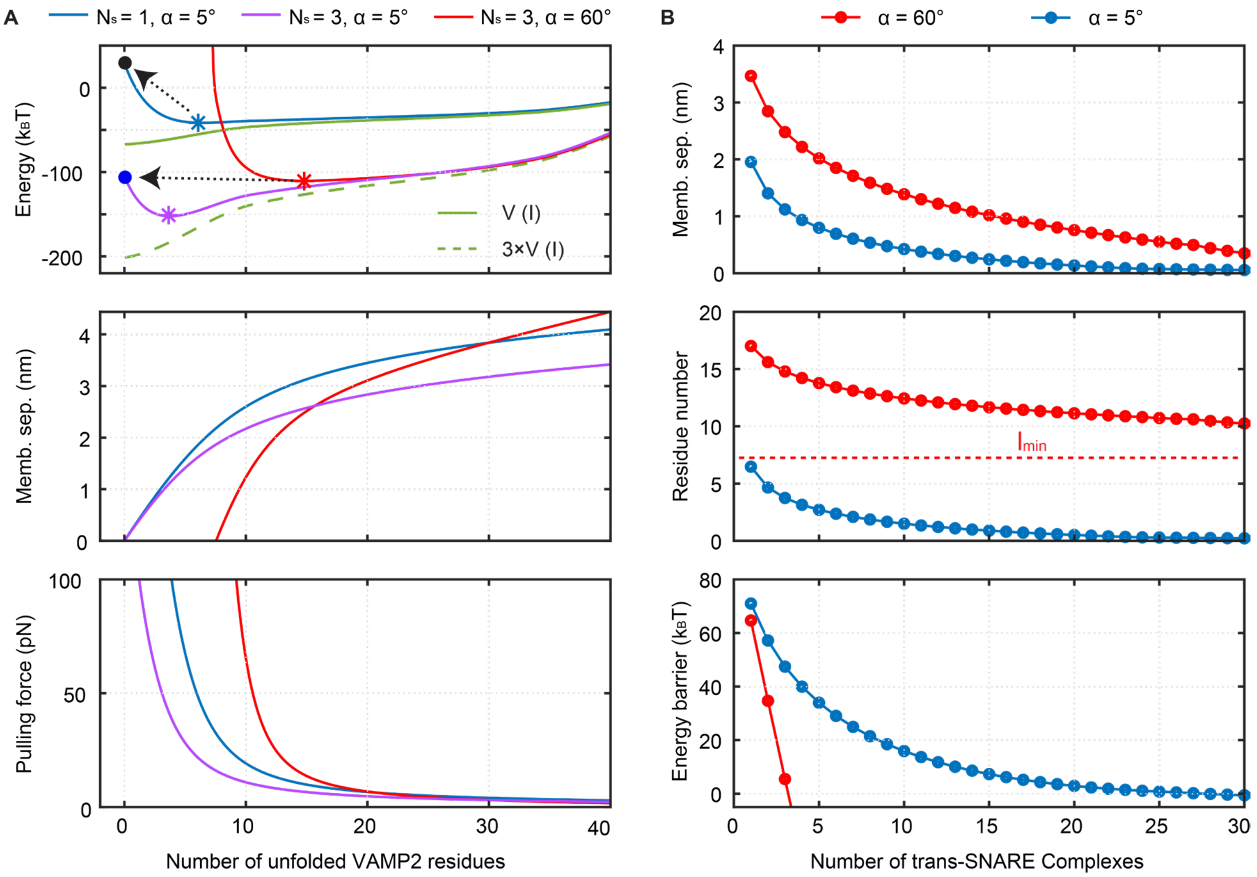


**Figure S18. A large syntaxin tilt angle promotes cooperative and productive assembly of multiple trans-SNARE complexes.**

(A) Total energy of the SNARE-membrane system (top), equilibrium membrane separation (middle), and pulling force per trans-SNARE complex (bottom) as functions of the number of unfolded VAMP2 residues in the presence of different syntaxin tilt angles (α) and numbers of trans-SNARE complexes (Ns). In the top panel, energy barriers for membrane fusion are indicated by arrows.

(B) Equilibrium membrane separation (top), number of unfolded VAMP2 residues (middle), and energy barrier for fusion (bottom) as functions of the number of trans-SNARE complexes at two syntaxin tilt angles. In the middle panel, the dashed line marks the minimum VAMP2 unfolding for a tilt angle of 60°.

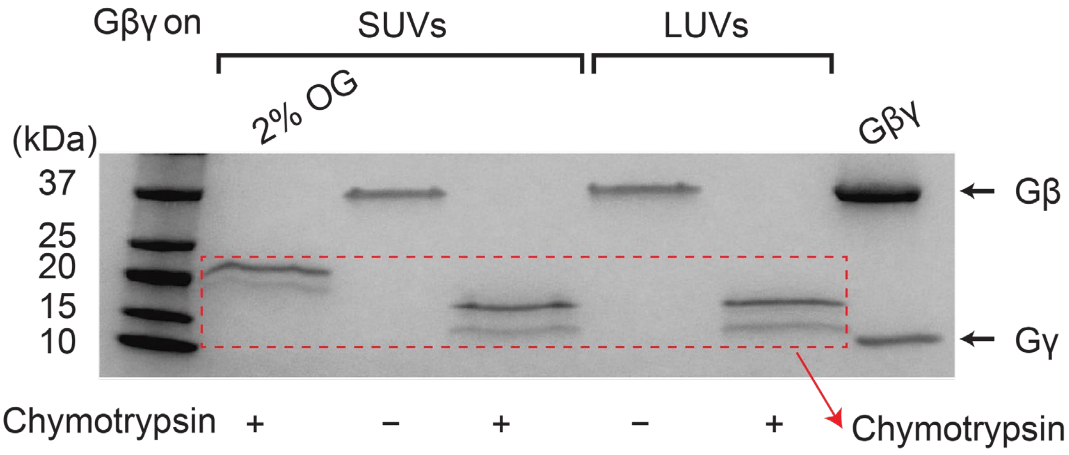


**Figure S19. Lipidated Gβγ heterodimers are located in the outer leaflet of vesicles.** Chymotrypsin digestion of lipid-anchored Gβγ complexes on SUVs or LUVs resulted in complete loss of both Gβγ subunits, demonstrating that Gβγ is exposed on the outer leaflet of the vesicle membrane. As a positive digestion control, 2% OG was first added to Gβγ-containing SUVs, and the sample was then treated with protease, leading to complete Gβγ digestion (second lane).

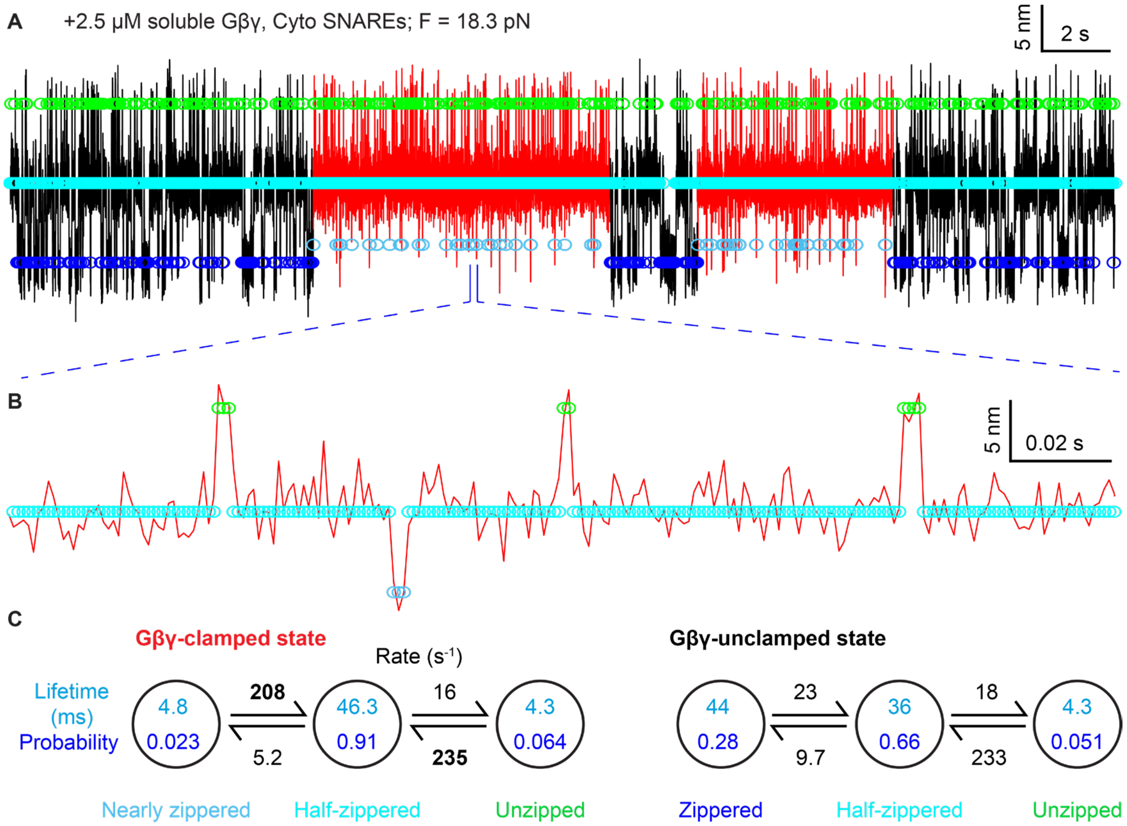


**Figure S20. Gβγ binds multiple SNARE conformations while clamping the half-zippered state.**

(A) Constant-mean-force extension trajectory, displayed at 1 kHz, corresponding to trace v in Figure 5C. The Gβγ-clamped region (red) and unclamped region (black) were analyzed separately with three-state hidden Markov models, yielding the state parameters shown in (C) and the idealized extensions indicated by colored circles: blue, zippered; light blue, Gβγ-bound nearly zippered; cyan, half-zippered; and green, unzipped. At this temporal resolution, the Gβγ clamp appears leaky, with brief excursions to a more zippered state whose mean extension is 1.9 nm greater than that of the fully zippered state, suggesting a highly dynamic Gβγ-clamped state.

(B) Expanded view of the indicated time interval in (A).

(C) State lifetimes, probabilities, and transition rates derived from hidden Markov modeling of the Gβγ-clamped and unclamped regions. Transition rates exceeding 200 $s^{-1}$ from the unzipped and nearly zippered states back to the Gβγ-clamped half-zippered state, shown in bold, suggest that Gβγ remained associated with the SNARE complex during these excursions. Complete dissociation followed by rebinding from 2.5 μM soluble Gβγ would require a bimolecular association rate constant of approximately $10^8$ M-1 s-1, near the upper range expected for productive protein–protein association.

**Table S1. SNARE constructs, membrane conditions, and pulling geometries used in this study.** Samples are grouped according to experimental configuration: cytosolic SNARE complexes; cis-SNARE complexes pulled approximately parallel to SUV or LUV membranes; variants testing lipid composition and bilayer geometry; membrane-anchored complexes pulled approximately perpendicular to SUVs; and trans-SNARE complexes bridging a t-SNARE-containing SUV and a VAMP2-containing nanodisc. For each sample, the table specifies the SNARE composition and engineered mutations or substitutions, lipid composition and bilayer geometry, and the direction through which force was applied. Unless otherwise indicated, complexes contained syntaxin-1A, VAMP2, and SNAP-25B and were crosslinked through engineered cysteines at the N-terminal −6 layer to permit reversible SNARE assembly. "WT" denotes the primary experimental constructs described in Supplementary Text T1, including cysteine substitutions introduced for site-specific crosslinking and tether formation, rather than completely native protein sequences. "Cyto" denotes constructs lacking the syntaxin and VAMP2 TMDs; "Syx" and "V2" denote syntaxin-1A and VAMP2, respectively; and "GS" denotes flexible glycine–serine sequences. LD, TMD, SUV, LUV, and $PIP_2$ denote linker domain, transmembrane domain, small unilamellar vesicle, large unilamellar vesicle, and phosphatidylinositol 4,5-bisphosphate, respectively.

| Sample name | SNARE proteins, mutations, or substitutions | Lipid or bilayer properties | Pulling geometry |
|---|---|---|---|
| Cyto | WT cytosolic complex; syntaxin-1A, VAMP2, and SNAP-25B; syntaxin and VAMP2 lack TMDs | No membrane | Cytosolic SNARE complex pulled through the C termini of syntaxin and VAMP2 |
| Cyto-LD2GS | Cytosolic complex; entire syntaxin LD (257–266) replaced with GSGGSGSSGS, and entire VAMP2 LD (85–94) replaced with GGSGSSGSGS | No membrane | Cytosolic pulling |
| CytoSyx_LD2GS | Cytosolic complex; entire syntaxin LD replaced by a GS linker; WT VAMP2 LD | No membrane | Cytosolic pulling |
| CytoSyx_LDC2GS | Cytosolic complex; syntaxin LD C-terminal residues 261–265 replaced with SGSSG; WT VAMP2 LD | No membrane | Cytosolic pulling |
| CytoV2_LD2GS | Cytosolic complex; entire VAMP2 LD replaced by a GS linker; WT syntaxin LD | No membrane | Cytosolic pulling |

| | | | |
|---|---|---|---|
| CytoV2_LDC2GS | Cytosolic complex; C-terminal five residues of the VAMP2 LD (90–94) replaced with SGSGS; WT syntaxin LD | No membrane | Cytosolic pulling |
| CytoV2_LDN2GS | Cytosolic complex; N-terminal portion of the VAMP2 LD (85–89) replaced with GGSGS; WT syntaxin LD | No membrane | Cytosolic pulling |
| **SNARE complexes pulled parallel to vesicle membranes** | | | |
| POPC-SUV | WT SNARE complex; syntaxin-1A, VAMP2, and SNAP-25B; syntaxin and VAMP2 contain TMDs | Pure POPC SUV; mean diameter 44 ± 13 nm | Cis-SNARE complex; both TMDs in the same SUV; pulled approximately parallel to the membrane |
| LD2GS | Entire LDs of both syntaxin and VAMP2 replaced by GS linkers | POPC SUV | Cis; parallel pulling |
| Syx_LDC2GS | C-terminal five residues of the syntaxin LD replaced by a GS linker; WT VAMP2 LD | POPC SUV | Cis; parallel pulling |
| V2_LD2GS | Entire VAMP2 LD replaced by a GS linker; WT syntaxin LD | POPC SUV | Cis; parallel pulling |
| V2_LDC2GS | C-terminal five residues of the VAMP2 LD replaced by a GS linker; WT syntaxin LD | POPC SUV | Cis; parallel pulling |
| LD-5GS-TMD | Five-residue GS linkers inserted between the LDs and TMDs of each protein: syntaxin, SGSSG; VAMP2, SGSGS | POPC SUV | Cis; parallel pulling |
| LD-10GS-TMD | Ten-residue GS linkers inserted between the LDs and TMDs of each protein: syntaxin, GSGGSGSSGS, VAMP2, GGSGSSGSGS | POPC SUV | Cis; parallel pulling |

| | | | |
|---|---|---|---|
| V2_TMD2PolyL | Entire VAMP2 TMD replaced by a polyleucine TMD; WT syntaxin LD | POPC SUV | Cis; parallel pulling |
| V2_TMD2PolyV | Entire VAMP2 TMD replaced by a polyvaline TMD; WT syntaxin LD | POPC SUV | Cis; parallel pulling |
| WT-TMD | Native TMD cysteines restored: syntaxin C272/C273 and VAMP2 C103 | POPC SUV | Cis; parallel pulling |
| **Lipid-composition and bilayer-geometry samples** | | | |
| 15%Cholesterol | WT SNARE complex | SUV: 85 mol% POPC and 15 mol% cholesterol | Cis; parallel pulling |
| 10%DOPS | WT SNARE complex | SUV: 90 mol% POPC and 10 mol% DOPS | Cis; parallel pulling |
| 2.5%$PIP_2$ | WT SNARE complex | SUV: 97.5 mol% POPC and 2.5 mol% $PIP_2$ | Cis; parallel pulling |
| Syx_LDC2GS–$PIP_2$ | C-terminal five residues of the syntaxin LD replaced by a GS linker; WT VAMP2 LD | SUV: 97.5 mol% POPC and 2.5 mol% $PIP_2$ | Cis; parallel pulling |
| V2_LDC2GS–$PIP_2$ | C-terminal five residues of the VAMP2 LD replaced by a GS linker; WT syntaxin LD | SUV: 97.5 mol% POPC and 2.5 mol% $PIP_2$ | Cis; parallel pulling |
| DMPC | WT SNARE complex | 100 mol% DMPC SUV; shorter 14:0/14:0 acyl chains and a thinner bilayer than POPC | Cis; parallel pulling |
| SAPC | WT SNARE complex | 100 mol% SAPC SUV; 18:0/20:4 acyl chains and a thicker bilayer than POPC | Cis; parallel pulling |
| POPC-LUV | WT SNARE complex | POPC LUV; mean diameter 135 ± 36 nm | Cis; parallel pulling |
| **Complexes pulled perpendicular to SUV membranes** | | | |
| Cyto-on-SUV | WT cytosolic complex; syntaxin attaches to | POPC SUV containing biotinylated lipids | Cytosolic complex positioned outside an SUV and pulled |

| | | | |
|---|---|---|---|
| | SUVs via biotin-streptavidin interaction | for DNA handle attachment | approximately perpendicular to the membrane; control for SUV deformation |
| Syx–CytoV2 | Full-length syntaxin anchored through its TMD; cytosolic VAMP2 lacking its TMD; SNAP-25B | POPC SUV | Cis-like complex anchored through syntaxin; pulled perpendicular to the membrane |
| CytoSyx–V2 | Cytosolic syntaxin lacking its TMD; full-length VAMP2 anchored through its TMD; SNAP-25B | POPC SUV | Cis-like complex anchored through VAMP2; pulled perpendicular to the membrane |
| Syx_LD2GS–CytoV2 | Full-length syntaxin with its entire LD replaced by GS and anchored through its TMD; cytosolic VAMP2; SNAP-25B | POPC SUV | Syntaxin-anchored complex pulled perpendicular to the membrane |
| Syx_LD-5GS-TMD–CytoV2 | Full-length syntaxin containing a five-residue GS insertion between its LD and TMD; cytosolic VAMP2; SNAP-25B | POPC SUV | Syntaxin-anchored complex pulled perpendicular to the membrane |
| **Trans-SNARE sample** | | | |
| Trans | Full-length syntaxin, VAMP2, and SNAP-25B | t-SNARE in a 44-nm SUV containing 99.5 mol% POPC and 0.5 mol% DSPE-PEG–biotin; VAMP2 in a 13-nm POPC nanodisc | Trans-SNARE complex bridging an SUV and a nanodisc |
| Trans-Syx_LD-5GS-TMD | Full-length syntaxin containing a five-residue GS insertion between its LD and TMD, SNAP-25B, and full-length VAMP2 | t-SNARE in a 44-nm SUV containing 99.5 mol% POPC and 0.5 mol% DSPE-PEG–biotin; VAMP2 in a 13-nm POPC nanodisc | Trans-SNARE complex bridging an SUV and a nanodisc |

### References


1. Stein, A., Weber, G., Wahl, M.C. & Jahn, R. Helical extension of the neuronal SNARE complex into the membrane. *Nature* **460**, 525-528 (2009).
2. Knecht, V. & Grubmuller, H. Mechanical coupling via the membrane fusion SNARE protein syntaxin 1A: A molecular dynamics study. *Biophys. J.* **84**, 1527-1547 (2003).
3. Li, F., Pincet, F., Perez, E., Eng, W.S., Melia, T.J., Rothman, J.E. & Tareste, D. Energetics and dynamics of SNAREpin folding across lipid bilayers. *Nat. Struct. Mol. Biol.* **14**, 890-896 (2007).
4. Kiessling, V., Kreutzberger, A.B., Liang, B.Y., Nyenhuis, S.B., Seelheim, P., Castle, J.D., Cafiso, D.S. & Tamm, L.K. A molecular mechanism for calcium-mediated synaptotagmin-triggered exocytosis. *Nat. Struct. Mol. Biol.* **25**, 911-917 (2018).
5. Marko, J.F. & Siggia, E.D. Stretching DNA. *Macromolecules* **28**, 8759-8770 (1995).
6. Keeble, A.H., Turkki, P., Stokes, S., Anuar, I.N.A.K., Rahikainen, R., Hytonen, V.P. & Howarth, M. Approaching infinite affinity through engineering of peptide-protein interaction. *Proc. Natl. Acad. Sci. USA* **116**, 26523-26533 (2019).
7. Guo, Z.L., Hong, H.Y., Sun, H., Zhang, X.F., Wu, C.X., Li, B., Cao, Y. & Chen, H. SpyTag/SpyCatcher tether as a fingerprint and force marker in single-molecule force spectroscopy experiments. *Nanoscale* **13**, 11262-11269 (2021).
8. Gao, Y., Zorman, S., Gundersen, G., Xi, Z.Q., Ma, L., Sirinakis, G., Rothman, J.E. & Zhang, Y.L. Single reconstituted neuronal SNARE complexes zipper in three distinct stages. *Science* **337**, 1340-1343 (2012).
9. Ma, L., Rebane, A.A., Yang, G., Xi, Z., Kang, Y., Gao, Y. & Zhang, Y.L. Munc18-1-regulated stage-wise SNARE assembly underlying synaptic exocytosis. *Elife* **4**, e09580 (2015).
10. Zhang, X.M., Rebane, A.A., Ma, L., Li, F., Jiao, J., Qu, H., Pincet, F., Rothman, J.E. & Zhang, Y.L. Stability, folding dynamics, and long-range conformational transition of the synaptic t-SNARE complex. *Proc. Natl. Acad. Sci. USA* **113**, E8031-E8040 (2016).
11. Rebane, A.A., Ma, L. & Zhang, Y.L. Structure-based derivation of protein folding intermediates and energies from optical tweezers. *Biophys. J.* **110**, 441-454 (2016).
12. Hernandez, J.M., Stein, A., Behrmann, E., Riedel, D., Cypionka, A., Farsi, Z., Walla, P.J., Raunser, S. & Jahn, R. Membrane fusion intermediates via directional and full assembly of the SNARE complex. *Science* **336**, 1581-1584 (2012).
